# RNA structural heterogeneity in a eukaryotic cell influences its heat shock response

**DOI:** 10.1101/2025.07.06.663400

**Authors:** Jiaxu Wang, Han Jian, Wen Ting Tan, Anthony Cheng, Jong Ghut Ashley Aw, Yue Wang, Guisheng Zeng, Niranjan Nagarajan, Yue Wan

## Abstract

RNA structure-based gene regulation remains under-explored in eukaryotes. While RNA can form different conformations in solution, the extent to which it folds into structure ensembles and how the different conformations regulate gene expression still needs to be fully understood. We coupled the SHAPE compound NAI-N3 with direct RNA sequencing to identify structure modifications along a single RNA molecule (sm-PORE-cupine). Using a combination of base mapping and direct signal alignment, we boosted the percentage of mappable RNA molecules from direct RNA sequencing. Using Bernoulli Mixture Model (BMM) clustering, we show that we can separate RNA structure ensembles from ligand-bound and unbound riboswitches accurately, identify isoform-specific structure ensembles along the SARS-CoV-2 genome, and determine RNA structure ensembles in the transcriptome of a eukaryote, *C. albicans*, at yeast (30°C) and hyphae (37°C) states. We observed that RNAs are more structurally homogenous at 37°C compared to 30°C, are more variable *in vivo* than *in vitro*, and show higher homogeneity in 3’UTRs than in the coding region. We also identified structure ensembles that are associated with changes in translation efficiency and decay in *C. albicans* at 30°C and 37°C and validated translational changes using reporter assays. Our work shows that single-molecule RNA structure probing using direct RNA sequencing can be applied to diverse transcriptomes to study the complexity and function of RNA structures.

## Introduction

While there are many examples of functional RNA structures in bacteria and viruses, how RNA structures could regulate gene expression in eukaryotic cells remains an important open question. As RNAs can fold into different conformations *in vitro* and *in vivo*, identifying functional RNA structures, how they change, and determining which structure in a population ensemble is biologically important enables us to understand fundamental RNA biology and how RNA can be targeted in diseases^1,2^.

To obtain single-molecule RNA structure information, several methods, including DRACO^3^, DREEM^4^, DANCE-MaP^5^, and Da Vinci^6^, have been developed to couple DMS or SHAPE chemical probing with high-throughput sequencing to read out RNA modifications along a single read of RNA. These modified patterns are then clustered using different algorithms to identify potential structure clusters that are present inside cells or in solution. Single-molecule RNA structure studies have identified structure ensembles along viral RNAs, such as in HIV^4^ and SARS-CoV-2 genomes^3,7^, human RNAs, such as in 7SK^5^, and plant RNAs, including in *COOLAIR*^6^, to better study functional RNA structures that could impact gene regulation. While useful, most of these strategies require the conversion of RNA into cDNA molecules and focus on a specific transcript of interest inside the cell.

We had previously coupled SHAPE chemical probing with nanopore direct RNA sequencing to determine aggregate RNA structure information on gene-linked RNA isoforms (PORE-cupine)^8^. Here, we further optimized SHAPE chemical probing and utilized signal alignments from direct RNA sequencing to identify RNA modifications along a single RNA molecule. We also tested different clustering strategies to group modified RNA reads into their corresponding structures. We named our single-molecule RNA structure probing method single-molecule PORE-cupine (sm-PORE-cupine). Using sm-PORE-cupine, we show that we can accurately separate riboswitch structure ensembles, identify RNA structure ensembles along different SARS-CoV-2 subgenomic RNAs (sgRNA), and determine structural heterogeneity in the *C. albicans* transcriptome. Our results demonstrate that RNAs are highly heterogeneous in transcriptomes and that changes in structural ensembles can influence gene regulation inside cells.

## Results

### High modification rates and low false positive rates enable accurate clustering of RNA structure populations

To identify paired and unpaired bases along an RNA, we had previously sequenced RNAs that are modified and unmodified with the SHAPE compound, NAI-N3^9^, using direct RNA sequencing^8^. We observed that NAI-N3-modified bases can shift the current mean and standard deviation during direct RNA sequencing, as compared to unmodified bases, to enable us to provide an aggregate structure signal across many molecules for a particular RNA. As the same RNA can fold into different conformations and direct RNA sequencing enables one molecule to thread through a pore at a time, we were curious whether we could separate structure populations based on their modification pattern per molecule (**Figure 1a**). As many factors could influence our ability to cluster RNA modification patterns along each molecule, we performed simulation experiments using synthetic reads to determine the impact of structural similarity, read depth, read length, and modification rate on our ability to form structural clusters accurately (**Supp. Figure 1a**). We observed that increasing the sequencing depth, length, and modification rate of the reads all improve our ability to separate structural populations, and that a modification rate of >1.5% coupled with longer read lengths of >750 bases can separate structure populations at a sequencing depth of 1000 reads/transcript (**Supp. Figure 1a**).

**Figure 1.**
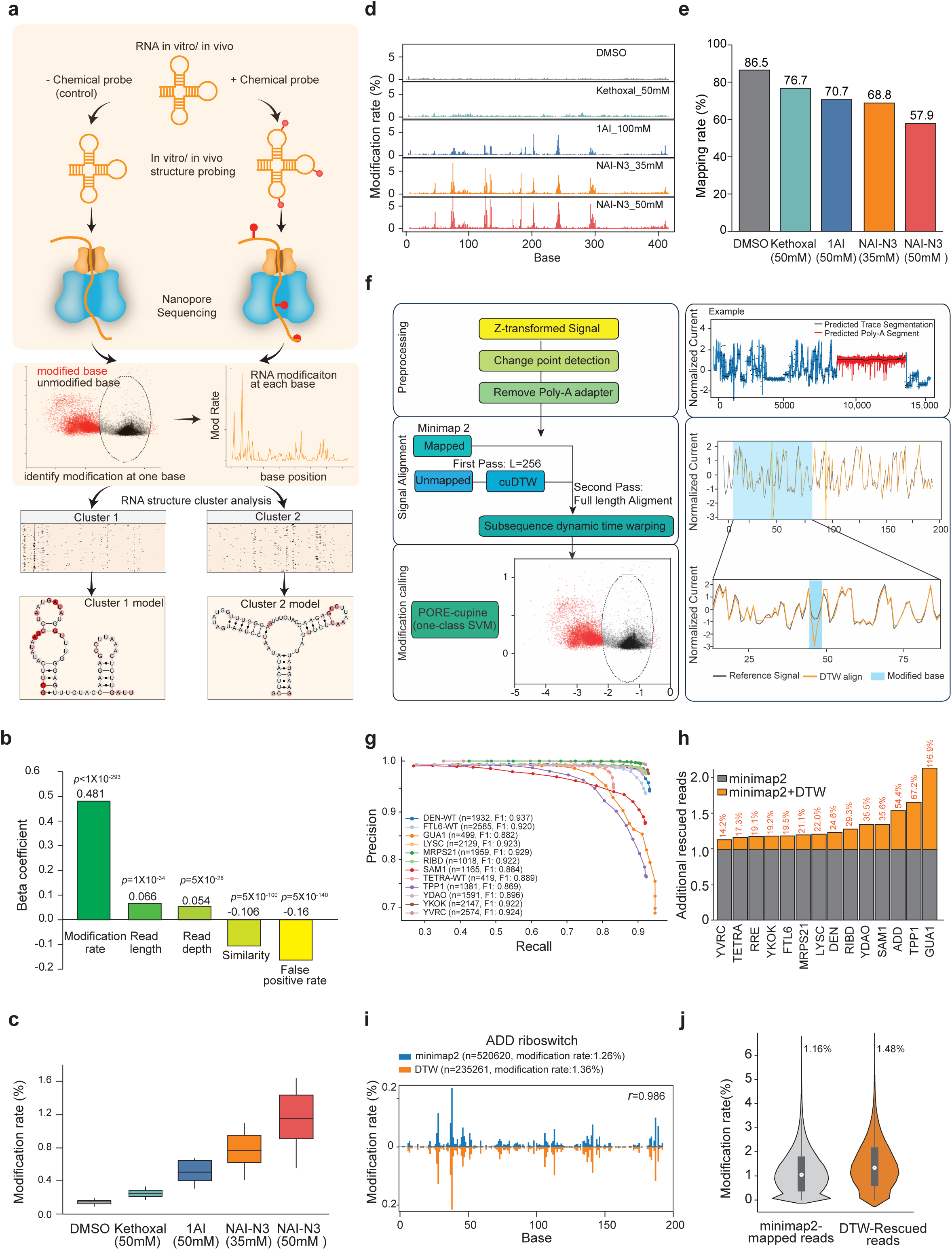
Experimental and analytical workflow of sm-PORE-cupine. **a,** Schematic of the experimental workflow to determine RNA structure ensembles from single-molecule RNA structure probing data using nanopore direct RNA sequencing. **b,** Bar chart shows the effects of each parameter (modification rate, read length, read depth, similarity, and false positive rate) in enabling accurate structure clustering, using a linear regression model on simulated data. **c**, Boxplot showing the distribution of modification rate using different chemical probes on a set of riboswitches and riboSNitches, and the modifications were identified by PORE-cupine^8^. **d**, Line plots showing the modification rate at each base after treating the Tetrahymena ribozyme with different chemical compounds. **e**, Bar chart showing the percentage of mapped reads by minimap2 for the different structure probing conditions tested. **f,** Left, Schematic showing the workflow for processing the sequencing signals, identifying the poly-A tail, and using sm-PORE-cupine to determine modifications. Right, we show an example of preprocessing (***Top***) and the signal alignment and modification calling on a modified site (***Bottom***). **g**, Precision-Recall curves for subsequence Dynamic Time Warping (subseqDTW) alignments with a truth set based on minimap2 alignments. F1 = (2 * precisions * recalls)/(precisions + recalls). **h**, Bar charts showing the percentage of reads that can be mapped for each RNA, using either minimap2 alone or minimap2 with DTW. The percentage of additional reads rescued by subseqDTW is in orange. **i**, Line plots showing the modification rates along the ADD riboswitch when the reads are mapped using minimap2 (blue) or DTW (orange). The Pearson correlation coefficient between the blue and orange plots is *r*=0.986. **j**, Violin plots showing the distribution of modification rates of reads that are mapped using minimap2 or rescued using DTW.

Additionally, as different ways of identifying SHAPE modifications using direct RNA sequencing, such as using PORE-cupine or TOMBO^10^, could have different sensitivities and false positive rates in detecting structure modifications, we also tested the effect of false positive rate (FPR) on our ability to separate synthetic RNA populations (**Supp Figure 1b**). We observed that high FPR significantly decreases our ability to cluster RNA structure populations. To determine the most important parameters out of all tested, we performed linear regression on them and identified that high modification rates and low false positive rates have the biggest impact on our ability to separate RNA structure populations accurately (**Figure 1b**).

### High concentrations of NAI-N3 can modify RNAs to high modification rates but are poorly sequenced and mapped by direct RNA sequencing

To increase the modification rate on RNAs for better separability, we were curious whether there are chemical probes that can detect and modify RNA structures with high efficiency and also be sequenced using direct RNA sequencing. As such, we tested several compounds, including kethoxal^11^, a smaller SHAPE compound 1AI, as well as lower and higher concentrations of NAI-N3, for their ability to modify a group of positive control RNAs and to be sequenced using direct RNA sequencing (**Figure 1c**). We observed that 50mM NAI-N3 resulted in the highest modification rate along the RNAs (**Figure 1c, d**) and that NAI-N3 modifications along an RNA can span a large distance along the RNA (**Supp. Figure 1c**), indicating that the modifications are not clustered together within a short distance. We also confirmed that a high concentration of NAI-N3 treatment resulted in highly accurate structure probing and that the modifications are found in single-stranded bases along an RNA, as expected (**Supp. Figure 1d**). As high modification rates along an RNA can result in blockages during direct RNA sequencing, decreasing the number of reads that can be sequenced and mapped properly, we also checked the number of sequencing reads from RNAs that are modified with kethoxal, 1AI, and low and high concentrations of NAI-N3. We observed that RNAs that are modified with high (50mM) NAI-N3 showed poorer mappability of sequencing reads (**Figure 1e**).

### Direct signal alignment using dynamic time warping can rescue >28% of failed reads

As the RNA threads through the pore during sequencing, each 5-mer has an associated current signature, resulting in a current signal that is unique to each RNA^12^. As dense modifications along an RNA can result in frequent signal deviations and thus poor base-calling and mappability using minimap2^13^, we investigated whether we could map the reads by directly aligning the current signals to the expected signals (**Figure 1f**). To test this, we performed dynamic time warping (DTW) based alignments on 14 benchmark RNAs that were treated with 50mM NAI-N3 (**Supp. Tables 1, 2**). We first checked that we could map the signals with high precision to the correct transcripts for both high and low quality reads (**Figure 1g, Supp. Figure 2a**).

Sequencing reads that are considered high quality upon sequencing are usually mapped with high efficiency, whereas low-quality sequencing reads are mapped inefficiently with minimap2. Even for high-quality reads, we observed that signal alignments can rescue >10% of these reads, increasing the percentage of high-quality mappable reads from 70.3% to 82.2% (**Supp. Figure 2b**). For low quality reads, which were traditionally mapped poorly by minimap2, we were able to increase the percentage of these reads that are mapped from 17.1% to 45.4%, increasing the total percentage of mappable reads from 56% to 73% for the 14 RNAs for downstream analysis (**Figure 1h, Supp. Figure 2b**). We observed that while other strategies for identifying modifications, such as using Tombo, can have a higher modification rate, DTW has the highest fold change of true modification rate over false positive rate (**Supp. Figure 2c, d**). We confirmed that NAI-N3 modifications detected from signal alignment are highly correlated with the modifications obtained from high-quality reads mapped with minimap2 (**Figure 1i, Supp. Figure 2e, 3a**), indicating that structure information obtained from direct signal alignments is accurate. Additionally, we confirmed that the failed-quality reads contain a higher modification rate than passed-quality reads and that failed reads that are rescued by DTW also have higher modification rates than reads that can be mapped by minimap2 (**Figure 1i, j, Supp. Figure 3a, b**). As a high modification rate is important for effective clustering, these rescued failed reads are valuable for downstream clustering of RNA structure populations.

### sm-PORE-cupine separates RNA structure populations of known riboswitches accurately

To determine the best way to cluster single molecules according to their reactivity profiles, we tested different clustering strategies, including BMM, Spectral^3^, DREEM^4^, and K-means^14,15^ clustering, using simulated data (**Methods**)^3,4^. We observed that BMM clustering generally achieves the highest accuracy in separating structural populations at a fixed sequencing depth and modification rate (**Supp. Figure 4a, b**), as well as for longer and shorter RNA molecules (**Supp. Figure 4c**). Additionally, we also observed that BMM clustering could be calculated at a faster speed than using DREEM (**Supp. Figure 4d**), suggesting that it could be particularly useful for transcriptome analysis. We proceeded to remove the bottom 25% (uninformative) and top outliers among reads in terms of modification rate, and to use BMM clustering for the remaining reads in our downstream analysis (**Supp. Figure 4e, Methods**).

To test our ability to separate real structure ensembles, we performed structure probing of the adenosine riboswitch in the presence and absence of adenosine (**Figure 2a**). We then mixed equal proportions of the two structure populations *in silico* and used BMM clustering to separate the single molecules into two clusters based on their reactivity profile per molecule. We observed that we could group the molecules into two clusters, with a majority of the molecules belonging to the No-ligand-bound riboswitch population in cluster 1 (73.7%) and a majority of the molecules belonging to the Ligand-bound riboswitch assay in cluster 2 (71.0%, **Figure 2b**). In addition to the adenosine riboswitch, we also confirmed that we can determine RNA structural ensembles from other Ligand-bound and No-ligand-bound riboswitches, including YKOK and Guanine (GUA1) riboswitches (**Supp. Figure 5a**-d). sm-PORE-cupine’s modification rate and clustering ability are also on par with the current gold standard for single-molecule RNA structure clustering using DMS-MaP-Seq and DREEM (**Figure 2b, Supp. Figure 5e**), and outperform PORE-cupine coupled and Tombo coupled with BMM (**Supp. Figure 5f**), confirming that we can group RNA structures accurately.

**Figure 2.**
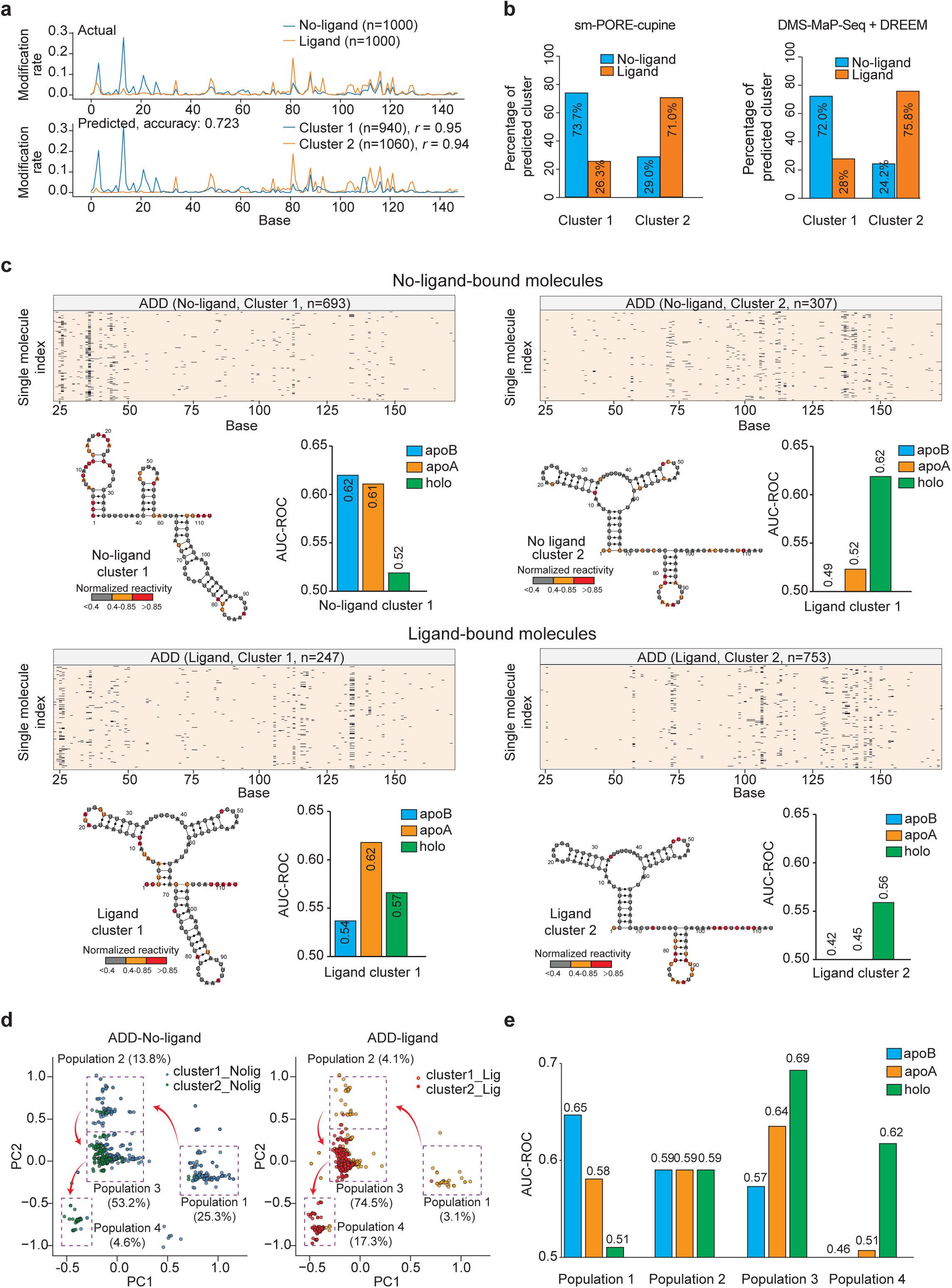
sm-PORE-cupine can identify structure populations in the ADD riboswitch. **a**, ***Top***, Line plot showing the aggregate reactivity of the ADD riboswitch when it is structure probed in the absence (blue) or presence (orange) of 5mM of adenosine. ***Bottom***, Line plot showing the aggregate reactivity from the two structure populations that are clustered using BMM (Methods). **b**, Bar charts showing the assignment of reads from Ligand-bound and No-ligand-bound ADD riboswitch into two structure populations based on sm-PORE-cupine with BMM clustering (***Left***) or DMS-MaP-Seq with DREEM clustering (***Right***). **c**, ***Top***, Heat map showing the distribution of NAI-N3 modifications along each molecule for molecules that are correctly (***Left***) or wrongly (***Right***) assigned as No-ligand bound by BMM clustering. The bar charts show the AUC-ROC calculated using the aggregate reactivity of the cluster against known structures of apoA, apoB, and holo. The aggregate reactivity of the cluster is then plotted on the known secondary structure with the highest AUC-ROC. ***Bottom***, Heat map showing the distribution of NAI-N3 modifications along each molecule for molecules that are correctly (***Right***) or wrongly (***Left***) assigned as Ligand-bound by BMM clustering. The bar charts show the AUC-ROC calculated using the aggregate reactivity of the cluster against known structures of apoA, apoB, and holo. The aggregate reactivity of the cluster is then plotted on the known secondary structure with the highest AUC-ROC. **d**, Principal component analysis showing the single molecule RNA structure trajectory for the ADD riboswitch in the absence and presence of adenosine. The proportion of each RNA population is shown. **e**, Bar plots showing the AUC-ROC values of each structure population in **d**, when assessed based on the apoA, apoB, and holo secondary structure model for the ADD riboswitch.

As the adenosine riboswitch can take on three different conformations corresponding to the No-ligand bound state (apoB), the Ligand-bound state (holo), and the intermediate state (apoA)^16^, we took a closer look at structures that are present in the different BMM clusters.

For the molecules that originated from the No-ligand-bound population, 693 molecules were classified correctly, whereas 307 molecules were misclassified (**Figure 2c**). To determine whether these molecules were truly misclassified or whether there are legitimate reasons behind this grouping, we plotted the reactivity along individual molecules for the “misclassified” and “correctly classified” molecules (**Figure 2c**), and calculated the accuracy of the aggregate reactivity of the clusters relative to the apoB, apoA and holo structures^16^. As expected, the AUC-ROC values for the correctly classified unbound molecules showed that they are most similar to the apoB state, although they also show high similarity to the intermediate apoA state (**Figure 2c**). Interestingly, we observed that a small “misclassified” No-ligand bound population resembles most closely the holo state, indicating that a major population of apoB or apoA structure and a minor population of holo structure conformation exist in the No-ligand bound population. For the Ligand-bound population, 753 and 247 of the molecules are classified correctly and incorrectly, respectively. The AUC-ROC values for the aggregate reactivity of the major cluster are the highest for the Ligand-bound structure (holo), although the smaller “misclassified” population shows higher similarity to the intermediate structure, suggesting that both the intermediate (apoA) and Ligand-bound structures (holo) exist in the Ligand-bound population (**Figure 2c**).

We additionally used principal component analysis (PCA) in ADD No-ligand and ADD Ligand samples based on the modification profile of each molecule to visualize this diversity of RNA conformations (**Figure 2d**). We observed that the number of molecules decreased in populations 1 and 2 and increased for populations 3 and 4 as we transition from the ADD No-ligand sample to the ADD Ligand sample. Interestingly, the AUC-ROC values based on the modification signal for population 1 are highest relative to the apoB conformation, while the AUC-ROC values increase from population 1 to population 3 and become very low in population 4 when compared to the apoA conformation. Finally, the AUC-ROC values increased from population 1 to population 3, and still remain high for population 4 when compared to the holo conformation (**Figure 2e**), highlighting the structural dynamic changes from no ligand to ligand binding for the ADD riboswitch.

Lastly, to test whether we can detect alternative structure populations when they are present in low proportions, we tested our ability to identify structure ensembles for a range of values for the presence of the alternative structure. We observed that we can identify structural ensembles even when only 10% of the alternative structure is present (**Supp. Figure 5g**), suggesting that our method is sensitive.

### The 3’end of the SARS-CoV-2 genome is highly heterogeneous

To determine the degree of heterogeneity in a region, we devised a homogeneity score that takes into account the degree of similarity of structural profiles between two clusters and the relative proportion of the clusters (**Methods**). To identify structural heterogeneity along a long RNA, we performed sm-PORE-cupine on a published NAI-N3-treated SARS-CoV-2 direct RNA sequencing dataset in Vero cells. As the presence of RNA structure ensembles can also show up as alternative RNA-RNA interactions in RNA proximity ligation experiments, we overlapped our heterogeneous regions with SPLASH data^17^. We observed that regions with many alternative pairwise RNA-RNA interactions (hubs) exhibit greater RNA structure heterogeneity than non-hub regions (**Supp. Figure 6a**), and that highly homogenous regions tend to have lower numbers of hubs (**Supp. Figure 6b**), providing orthogonal evidence that our measured homogeneity score is meaningful. Interestingly, we observed that the 3’ end of the SARS-CoV-2 genome is highly heterogeneous as compared to the rest of the genome based on both sm-PORE-cupine data and SPLASH interaction hubs (**Supp. Figure 6c**).

The 3’end of the SARS-CoV-2 genome contains subgenomic RNAs that are generated by discontinuous transcription^18^. We had previously shown that different subgenomic RNAs, including RNAs that encode ORF7a, ORF8, and nucleocapsid (N), can fold into different conformations in cells^17^. As such, the increased heterogeneity observed at the 3’ end of the SARS-CoV-2 genome could be due to the presence of different RNA structures of different subgenomic RNAs. To determine whether we can identify heterogeneous regions along each sgRNA, we identified and grouped reads that belong to each full-length subgenomic RNA. Out of the different subgenomic RNAs, we obtained sufficient coverage for N, ORF7a, and ORF8 for RNA heterogeneity analysis (**Supp. Figure 6d**). We used the same number of sequencing reads from each RNA for heterogeneity analysis and observed that N is the most heterogeneous out of the three (**Supp. Figure 6e**). This suggests that the same sequence along different subgenomic RNAs can fold more readily into different ensembles. To highlight this complexity, we examined a region in the 3’UTR that was previously reported to form two conformations. Using the same number of sequencing reads across N, ORF7a, and ORF8 sgRNA, we observed that this region’s homogeneity score lies in the bottom 20% of SARS-CoV-2 windows, suggesting that it folds into at least two structural populations (**Supp. Figure 7a**). However, the relative proportions of the two conformations vary in different subgenomic RNAs (**Supp. Figure 7b, c**), even in regions with the same sequence.

### sm-PORE-cupine identifies heterogeneous regions in the *C. albicans* transcriptome

As most of the current RNA structure ensemble studies have been performed on individual transcripts, we applied sm-PORE-cupine to study RNA structural heterogeneity in the transcriptome of the eukaryote *C. albicans*. As *C. albicans* undergoes yeast-to-hyphae transition when the temperature increases from 30°C to 37°C^19^, we wanted to determine whether any RNA structural elements could regulate translation during this transition (**Figure 3a**).

**Figure 3.**
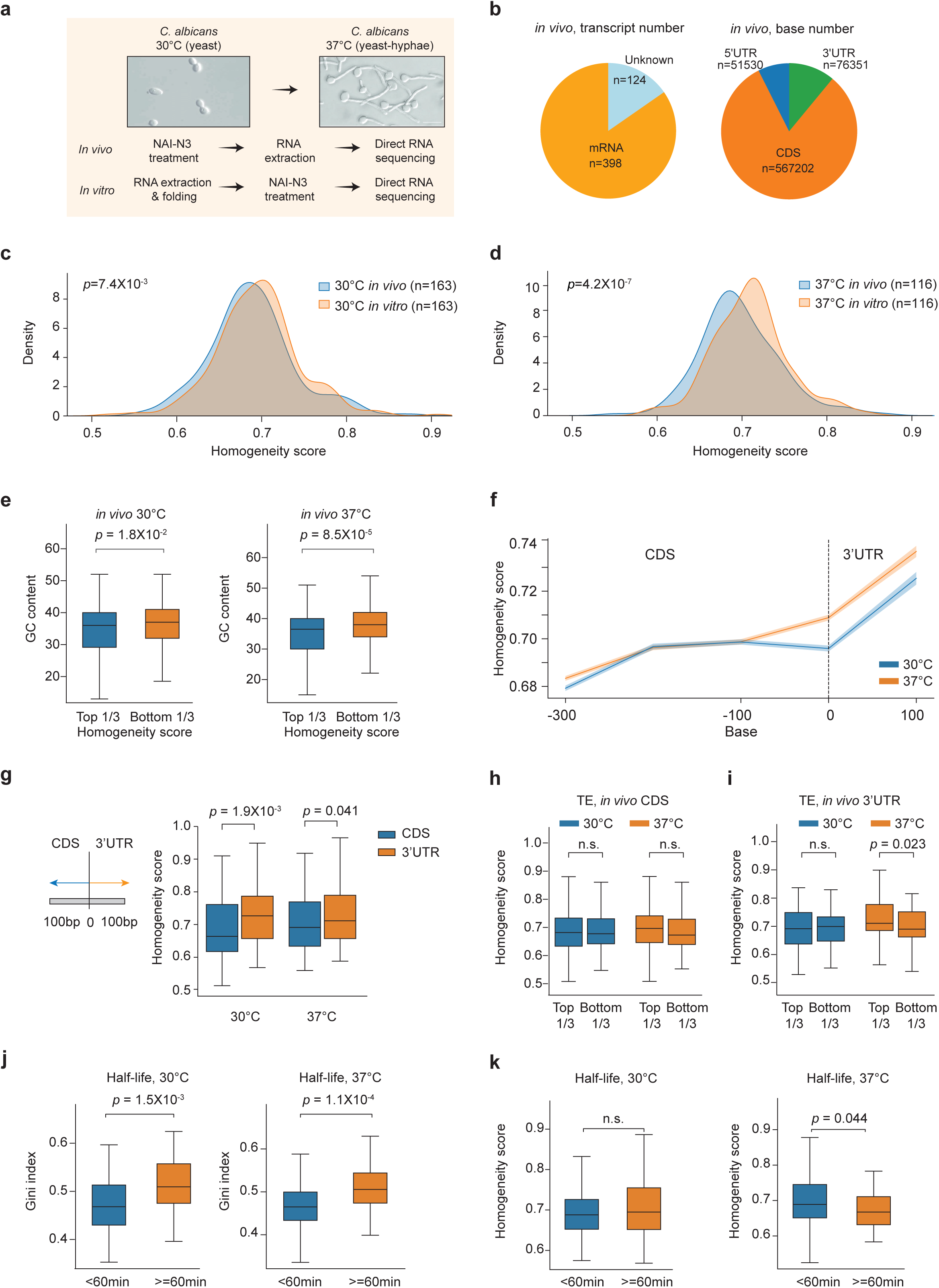
sm-PORE-cupine finds structural populations and their features in the *C. albicans* transcriptome. **a**, Images of *C. albicans* grown at 30°C (yeast) and 37°C (hyphae) respectively. **b**, Pie chart showing the number of detected transcripts (***Left***) and bases (***Right***) in *C. albicans* that we can perform sm-PORE-cupine on. **c, d**, Density plots showing the distribution of structural homogeneity in the *C. albicans* transcriptome when the structure probing is performed *in vivo* versus using refolded RNA *in vitro*, at 30°C (**c**) and 37°C (**d**). *P*-values are calculated using the two-sample Kolmogorov-Smirnov test. **e,** Boxplots showing the distribution of GC content in windows with high or low homogeneity scores *in vivo* at 30°C (***Left***) and 37°C (***Right***). *P*-values are calculated using the Wilcoxon Rank-Sum test. **f,** Metagene analysis of structural heterogeneity of *C. albicans* mRNAs at CDS and 3’UTR in 30 °C (blue) and 37 °C (orange). **g,** Boxplots showing the distribution of homogeneity scores in the last 100nt window of the CDS and the first 100nt window of the 3’UTR, at 30°C and 37°C. *P*-values are calculated using the Wilcoxon Rank-Sum test. **h, i**, Boxplots showing the distribution of homogeneity scores in CDS (**h**) or 3’UTR (**i**) for transcripts with high (top 1/3) or low (bottom 1/3) translation efficiency at 30 °C or 37 °C. *P*-values are calculated using the Wilcoxon Rank-Sum test. **j**, Boxplots showing the distribution of Gini index in transcripts that are degraded under 60min and in transcripts that have a longer half-life >=60min, at 30°C and 37°C. *P*-values are calculated using the Wilcoxon Rank-Sum test. **k**, Boxplots showing the distribution of homogeneity score in transcripts that have shorter half-lives(<60min) or longer half-lives (>=60min), at 30°C and 37°C. *P*-values are calculated using the Wilcoxon Rank-Sum test.

We performed two biological replicates of structure probing on *C. albicans* at 30°C and 37°C, both *in vivo* and *in vitro*. In total, we sequenced 18.1 million reads at 30 °C and 17.8 million reads at 37 °C, respectively, with a median length of 297-465 bases for the reads (**Supp. Figure 8a, b, Supp. Table 2**). We optimized signal alignment efficiency to enable faster mapping for whole transcriptome with high precision (**Supp. Figure 8c**), and observed that the signal alignments enabled us to rescue an additional 22.6% of reads for downstream analysis (**Supp. Figure 8d**). Using both reads aligned by DTW and minimap2, we observed a high correlation between the two biological replicates (*R*>0.83, **Supp. Figure 8e, f**), demonstrating the robustness of our assay.

We captured RNA structure information for 398 and 282 transcripts *in vivo* and *in vitro,* respectively, including 5’UTR, 3’UTRs, and CDS regions (**Figure 3b, Supp. Figure 8g**). Scanning along the *C. albicans* transcriptome, we observed that RNA regions are more homogeneous when they are folded *in vitro*, as compared to *in vivo* (**Figure 3c, d**) and that highly heterogeneous regions are enriched for higher GC content both *in vivo* and *in vitro* (**Figure 3e**). Across the transcriptome, we observed that CDS regions were more heterogeneous than 3’UTRs (**Figure 3f, g**). We hypothesized that this increase in heterogeneity might be due to the CDS regions being translated and unwound by ribosomes. As such, we binned transcripts based on their high or low translation efficiencies in a published dataset^19^ and plotted the distribution of heterogeneous regions in both CDS regions and 3’UTRs. Interestingly, we did not observe a significant difference in heterogeneity in the CDS regions between highly and poorly translated mRNAs (**Figure 3h**), although we did observe that the 3’UTRs of highly translated transcripts tend to be more homogeneous than those of poorly translated transcripts at 37°C (**Figure 3i**), suggesting that the structural heterogeneity of 3’UTRs could play a role in translation.

In addition to translation, we also measured mRNA decay rates at 30°C and 37°C by performing 2 biological replicates of RNA sequencing at 0, 20, 60, and 120min after treating *C. albicans* with Thiolutin (**Supp. Figure 9a, Methods**). We then used the program, Stepminer^20^, to obtain decay information for mRNAs at 30°C and 37°C (**Supp. Figure 9b, c, Supp. Table 3, Methods**). We observed that mRNAs were generally degraded faster at 37°C than at 30°C (**Supp. Figure 9d, Supp. Table 3**). Transcripts that are degraded faster tend to have lower Gini index and higher RNA modification rate, suggesting that they are more single-stranded (**Figure 3j, Supp. Figure 9e**). Additionally, we observed that faster decay at 37°C is associated with increased structure homogeneity (**Figure 3k**).

### RNA structures tend to become more homogenous upon heat shock

Interestingly, when *C. albicans* was shifted from 30°C to 37°C, we observed that there is a subtle but significant global shift towards mRNAs becoming more structurally homogenous (**Supp. Figure 10a, b**). We identified a total of 95 regions in 3’UTRs that showed heterogeneity changes when the temperature was shifted from 30°C to 37°C **(Figure 4a, Supp. Table 4**). To study the RNA structure profiles more closely, we selected two RNAs, RPS19A and RPL29, that showed changes in 3’UTR heterogeneity. We clustered single-molecule structure profiles of these two mRNAs (**Figure 4b, c**), and also performed RNA structure modelling of the clusters to visualize the shifts in RNA secondary structures at higher temperatures **(Figure 4d, Supp. Figure 10c**). We confirmed that these 3’UTR regions indeed result in changes in translation efficiency by cloning them behind the firefly luciferase and measuring the amount of protein production at 30 °C and 37 °C *in vitro* (**Figure 4e, f**), agreeing with the published translation efficiency changes from 30 °C to 37 °C in vivo^19^(**Figure 4g**), suggesting these RNA regions could potentially serve as RNA thermometers to regulate protein production in *C. albicans*.

**Figure 4.**
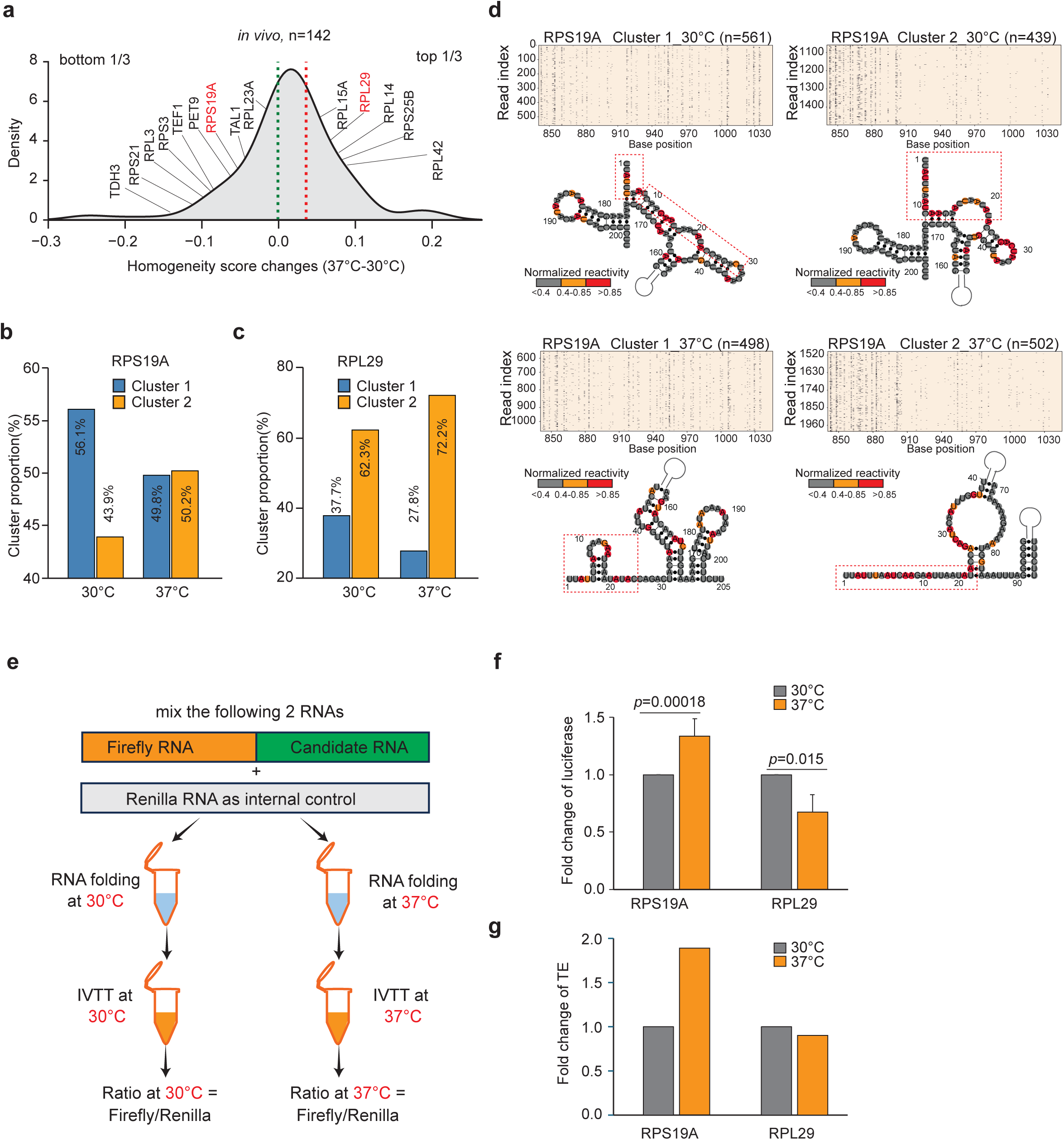
Structural populations in the 3’UTR of *C. albicans* transcripts can impact translation during heat shock. **a**, Density plot showing the distribution of RNA homogeneity changes between 30°C and 37°C. The selected transcripts for validation are in red. **b, c**, Barplots showing the distribution of RPS19A (**b**) and RPL29 (**c**) molecules in two different structural populations, at 30°C and 37°C. **d**, ***Top***, Heat map showing the distribution of NAI-N3 modifications along each molecule in each cluster of RPS19A using BMM. ***Bottom***, structure modelling was performed for each cluster using the aggregated reactivity as constraints. The red dash boxes indicate the same base positions along each secondary structure. High, medium, and low reactivity bases are shown in red, orange, and grey, respectively. **e**, Schematic showing the workflow to test the amount of protein production of a target 3’UTR, using *in vitro* transcription and translation assay. **f**, Bar plots showing the fold change in luciferase (***Top***) and in published translation efficiency data^19^ of target genes upon the temperature change from 30°C to 37 °C (***Bottom***).

## Discussion

Understanding how RNA structure impacts gene regulation and identifying functional structure populations that an RNA can adopt *in vivo* are key questions in the RNA structure field. While RNA-based gene regulation is important in eukaryotes, specific examples of RNA structure-based regulation remain rare. Here, we developed a new method (sm-PORE-cupine), which utilizes direct RNA sequencing to identify RNA structure heterogeneity and applied it to the SARS-CoV-2 genome and the transcriptome of the pathogenic fungi *C. albicans*. While previous structure ensemble studies typically focused on an RNA of interest, by optimizing the increased structure modification signal and accelerating signal alignments, we are now able to perform transcriptome-wide structure ensemble analysis in *C. albicans*. This helped us to better understand RNA structural properties *in vivo* and identify structure ensembles that could change and impact mRNA translation in *C. albicans*.

However, challenges in coupling chemical compounds such as SHAPE with direct RNA sequencing remain. These include titrating the amount of SHAPE compounds used such that RNAs are modified sufficiently and yet not stuck in the pores during sequencing. Additionally, 2’/3’ acetylation by SHAPE compounds can also result in 3’ end blockage, preventing adaptor ligation during the library preparation step. As each batch of NAI-N3 generated could have slight variations in their reaction efficiencies, depending on their source and how they have been stored, it is important to check the modification rate of each new batch of NAI-N3 generated by footprinting or dot blot experiments.

While we have previously identified the consensus RNA structures of SARS-CoV-2 subgenomic RNAs, our ability to identify structural ensembles in individual subgenomic RNAs provides an additional layer of resolution that is missing in the literature. As different subgenomic RNAs can fold differently in contributing to structure ensembles, these conformations could play a role in regulating gene expression for each subgenomic RNA. We anticipate that additional structure-function experiments will enable us to better understand how the different ensembles impact virus function.

Studying how RNAs fold *in vivo* and *in vitro* can enable our understanding of how RNAs are regulated by the cellular environment *in vivo.* Interestingly, we observed that RNA structures are more heterogeneous *in vivo*, as compared to *in vitro*, suggesting that some of the structures might be misfolded into kinetically trapped states i*n vitro*^21^. Additionally, concordant with another recent publication^22^, we also observed that RNA structures are more homogenous at higher temperatures both *in vivo* and *in vitro*, although the effect is stronger in vitro, suggesting that the melting of RNA structures could enable greater structural uniformity at higher temperatures and that there could be additional regulation in vivo to prevent large structural changes under heat shock. More experiments are needed to better dissect which functional structure, out of all possible conformations, could impact gene regulation at different temperatures.

While other methods, including coupling SHAPE with PacBio sequencing, can also enable single-molecule RNA structure ensemble determination^6^, we believe that the potential ability of direct RNA sequencing to detect both structure-probing-based and natural modifications in the future will enable deeper insights into the impact of RNA structure and modifications on gene regulation^23,24^. As direct RNA sequencing continues to improve in read throughput of untreated RNAs, we have checked that new version RNA004 for NAI-N3 treated RNAs, we observed 20 times more sequencing depth in RNA004 than RNA002 (**Supp. Figure 11a**), meanwhile, RNA004 can detect RNA structure modifications and result in clustering as well or better than the current RNA002 version (**Supp. Figure 11b, c**). We believe this approach will become more attractive and widely applied to study structure-function relationships across diverse organisms and biological systems.

## Methods

### Structure folding for riboswitches and riboSNitches

For structure mapping of the RNA benchmarks *in vitro*, we PCR amplified DNA fragments that contain the T7 promoter upstream of the RNA of interest. The DNA template was then *in vitro* transcribed (IVT) using NEB HiScribe Kit following the manufacturer’s instructions to generate the RNA. Briefly, we performed RNA structure probing as such: We diluted 1μg of each IVT RNA in water to a total volume of 16μl and folded the RNA in a PCR machine. The folding conditions are: we heated the RNA at 90°C for 2 min and chilled it on ice for 2 min, before adding 4μl of 5x structure probing buffer (500mM Tris pH 7.4, 50mM MgCl_2_, and 750 mM NaCl) to the mixture. We then ramped up the temperature of the PCR machine to 37°C at a rate of 0.1°C/s, and incubated the RNA for 20 min after the temperature was stabilized at 37°C. For structure probing in the presence of ligands, we added ligand or water control to the folded RNAs at 37°C and incubated for another 20 minutes. We then added NAI-N3 or other structure compounds as indicated in each figure and incubated for another 10 minutes. DMSO was used as a negative control for structure probing. After structure probing, the NAI-N3 and DMSO-treated RNA was purified using phenol: chloroform: isoamyl alcohol (25:24:1). The concentrations of NAI-N3 and other structure probing compounds were as shown in each figure. The working concentrations of the ligands for the riboswitches are: 5mM of adenosine for the adenosine riboswitch, 75μM of guanine for the GUA1 riboswitch, and 10mM of magnesium for the YKOK riboswitch.

### Direct RNA sequencing with Oxford Nanopore SQK-RNA002 kit on MinION systems

*C. albicans* RNA or IVT RNA preparation for Nanopore sequencing: 1μg of mRNA isolated from *C. albicans* sc5314 total RNA or in vitro transcribed (IVT) RNA was used for library preparation following the manufacturer’s protocol for the direct RNA library preparation kit SQK-RNA002 from Oxford Nanopore Technologies. *C. albicans* mRNA was isolated using the Poly(A) Purist MAG Kit (Thermo Fisher Scientific, AM1922).

RT Adapter ligation reaction and Reverse Transcription. To perform the first adapter ligation, we added 3μl of NEBNext Quick Ligation Reaction Buffer (NEB B6058), 0.25μl of RNA CS (RCS, 110nM), 1μl of RT Adapter (RTA) and 1.5μl T4 DNA Ligase (NEB, M0202) to 1μg of mRNA isolated from *C. albicans* sc5314 total RNA or to the in vitro transcribed (IVT) RNA in a total volume of 15μl. This reaction was incubated at room temperature for 10 min.

We then performed reverse transcription by adding 9μl of nuclease free water, 2μl of 10mM dNTPs (Thermo Scientific, R0192), 8μl of 5x first-strand buffer, 4μl of 0.1M DTT and 2μl of SuperScriptIII reverse transcriptase (Invitrogen 18080085) to the reaction and incubating at 50°C for 50 min, followed by 70°C for 10 min, and hold at 4°C.

We removed excess adapters by using RNAClean XP beads (Beckman Coulter, A63987), using a 1.8x beads ratio (using 72μl of RNAClean XP beads for a 40μl reverse transcription reaction). After binding the adapter-ligated RNA to the beads at room temperature for 5min, the beads were washed with 150μl of 80% ethanol. The reverse-transcribed RNA was then eluted in 23μl of nuclease-free water.

For the second adapter ligation, we added 8μl of NEBNext Quick Ligation Reaction Buffer, 6μl of RNA Adapter (RMX), and 3μl T4 DNA Ligase to 23μl of reverse-transcribed RNA in a total volume of 40μl. This reaction was incubated at room temperature for 10 min. We then removed the excess adapters by performing one round of RNAClean XP beads purification with a 1.0x beads ratio (using 40μl of RNAClean XP beads for a 40μl adapter ligation reaction). After binding the adapter-ligated RNA to the beads at room temperature for 5min, we washed the beads twice with 150μl of the Wash Buffer (WSB). The adapter-ligated RNA was eluted in 21μl of Elution Buffer (EB). The obtained RNA was then measured using a Qubit fluorometer DNA HS assay.

### Priming and loading of the SpotON flow cell

We primed and loaded the SpotON flow cell (R9.4.1) by following the manufacturer’s protocol. To prepare the flow cell priming mix, we thawed and mixed 30μl of Flush Tether (FLT) and added it directly to a tube of thawed and mixed Flush Buffer (FB) and mixed the solution by vortexing at room temperature. We first loaded 800μl of the priming mix into the flow cell via the priming port, and then loaded another 200μl of the priming mix into the flow cell after 5 min. We then added 17.5μl of nuclease-free water and 37.5μl of RNA Running Buffer (RRB) to the prepared 21μl of library sample. We loaded 75μl of sample into the Flow Cell via the SpotON sample port in a dropwise fashion before starting the sequencing run.

### Direct RNA sequencing with Oxford Nanopore SQK-RNA004 kit on PromethION

We used 600ng of in vitro transcribed (IVT) RNA for library preparation following the manufacturer’s protocol for the direct RNA library preparation kit SQK-RNA004 from Oxford Nanopore Technologies. The libraries were loaded onto FLO-PRO004RA PromethION RNA flow cells and sequenced on a PromethION 2 Solo device.

The library preparation steps were the same as those for SQK-RNA002 except the following: 1) 1μl of SUPERase·In™ RNase Inhibitor (Invitrogen, AM2694) is added during the RT Adapter ligation reaction, 2) 6μl of RNA Ligation (RLA) instead of 6μl of RNA Adapter (RMX) is added in the second adapter ligation reaction, 3) the adapter-ligated RNA is eluted in 33μl of RNA Elution Buffer (REB), and 4) we prepared the sample for loading by adding 100μl of Sequencing Buffer (SB) and 68μl of Library Solution (LIS) to 32μl of the library sample.

### *C. albicans* culture and hyphae induction

*C. albicans* (SC5314) was inoculated in 10ml YPD medium for 16-18 hours at 30°C, with constant shaking at 250rpm. We then transferred 2ml of the overnight culture to a fresh tube (1:50) of 100ml YPD medium with shaking for 3-5hours at 250rpm until the culture reaches an OD600 of 0.5-0.6. For hyphae induction, we added 10% FBS to the 30°C *C. albicans* culture, transferred it to 37°C with shaking at 250rpm for 1hour. For *in vivo* structure probing, we added NAI-N3 to a final concentration of 100mM with shaking of the culture for 10min. We then collected the cell pellet after centrifugation at 4000 RPM for 5min. The cell pellet was stored at −80°C until needed.

### Extraction of the *C. albicans* total RNA

To extract RNA from *C. albicans*, we first thawed the frozen pellet at room temperature and then placed it on ice. The pellets were re-suspended in 2.8ml yeast lysis buffer (100ml lysis buffer: 0.7ml 1M Tris-HCl, pH 7.0, 0.3ml 1M Tris-HCl, pH 8.0, 2ml of 0.5M EDTA, 5ml of 10%SDS). We transferred 700μl of this mixture into a 1.5ml Eppendorf tube, added 700μl of acid phenol, and vortexed rigorously. We then incubated the tubes at 65°C in a thermomixer for one hour, with constant shaking at 1000 rpm. After the incubation, we centrifuged the sample at maximum speed for 10 minutes at 4°C. We then transferred the aqueous phase into a new tube, added the same volume of fresh acid phenol, vortexed it, and spun it at maximum speed for 10 minutes at 4°C. We performed acid phenol extraction at least 2-3 times until the aqueous phase looked clear. To precipitate the RNA from the aqueous phase, we transferred the aqueous phase into a 2 ml tube and added 1/10 volume of 3M sodium acetate and 3x ice-cold 100% ethanol. We precipitated the RNA at −20°C for 1 hour before spinning it at maximum speed for 20-30 mins at 4°C. The pelleted RNA was then washed in 800μl of ice-cold 70% ethanol, centrifuged to remove the ethanol, and then air dried for 2 min before being dissolved in 50μl of nuclease-free water.

### IVTT and Luciferase Assay

In vitro translation reactions were prepared using Retic Lysate IVT Kit (Invitrogen, AM1200), and the luciferase assays were performed using Dual-Luciferase Reporter Assay System (#E1980). To set up the in vitro translation reaction, we mixed 150ng of the folded IVT Firefly RNA with 3’UTR targets, 120ng of Renilla IVT RNA, 0.5μl of 1:1 mix of –met and –leu 20x Translation Mix, 6.8μl of Retic lysate in a final volume of 10μl. The reaction mix was incubated at either 30°C or 37°C for 30 min. The in vitro translation reaction was then put on ice, and a 5μl aliquot of the reaction was added to 75 μl of 1x Passive Lysis Buffer. 20μl of the sample in Passive Lysis Buffer was then added to 100 μl of LARII in a 96-well plate, and the firefly luminescence was measured using a luminometer at an Integration of 1 second. 100μl of 1x Stop & Glo substrate in buffer was then added to each well, and the Renilla luciferase luminescence was measured by a luminometer at an Integration of 1 second.

### Measuring RNA decay in *C. albicans*

The RNA half-life was measured as previously described^25^. Briefly, *C. albicans* (sc5314) was inoculated into 10ml of liquid YPD media and grown overnight shaking at 30°C and 250rpm. Aliquot the overnight cultured Candida albicans cells to fresh medium (1:100), and continue culture at 30°C, 250rpm for around 3 hours until OD600=0.6. The aliquot Candida albicans culture was then centrifuged and resuspended in 30°C or 37°C prewarmed YPD medium. The Thiolutin (Merck, CAS RN 87-11-6) was then added to each aliquot to a final concentration of 10μg/ml. The culture aliquots were then cultured for 20min, 40min, 60min, and 120min at 30°C or 37°C, respectively. The cells were collected, and the RNA was extracted as described above, and the RNA library was then prepared using Illumina Nextra library prep kit. The library is sequenced on a Novaseq X to generate 2X150 base reads.

### Generation of simulated datasets

Synthetic datasets were generated to determine the optimal strategy for accurate clustering of RNA structure populations. To test the effect of structural similarity, RNA modification rate and sequencing depth on clustering, these parameters were varied in the simulations. The simulations begin by generating two template RNA structures (representatives of the two populations of RNA that we will simulate sequencing for) that have patterns of 0 and 1 corresponding to whether a position is double or single-stranded. The first template was generated with a binomial distribution defining the probability of being single-stranded at each position. A second synthetic template was then generated that has a defined average similarity (50% to 90%) to the template in terms of its modification pattern. For each of these templates, we then simulated RNA sequencing with modification rates varying from 0.1% to 10%, and sequencing depth varying from 1000 to 10,000 reads. False-positive modifications were also introduced into each simulated read based on a defined false-positive rate and an independent and identically distributed binomial model for each position.

### Clustering structure populations using a Bernoulli mixture model (BMM)

We adopted the Bernoulli mixture model (BMM) approach proposed in the DREEM paper^4^ with modifications to account for numerical stability and processing efficiency issues. BMM can cluster read data from alternative structures in RNA structure ensembles based on the assumption that: (1) each base of the read has a distinct mutation probability due to the probing efficiency of a compound, and (2) that base mutation probabilities are independent for each position on a read. The modifications in our BMM implementation include: (a) log transformation of the joint probability of observing a read **x**_n_ from cluster k in the applied Bernoulli mixture model to enable numerically stable calculations of small probabilities, and (b) use of a just-in-time compiler for Python to compile annotated code into machine code and significantly improve the computational efficiency of the implementation.

### Assessment of clustering accuracy and contributing factors

Clustering accuracy was assessed based on the percentage of reads that are assigned correctly to the most similar cluster based on ground truth labels (i.e. this translates to [<dominant label % in cluster 1> + <dominant label % in cluster 2>]/2, where distinct labels are assigned to the two clusters). BMM clustering was only performed with informative reads with modification rates between 0.75%-2.5% to avoid the impact of noisy reads on clustering. A linear regression model was used to assess the impact of parameters such as structural similarity, read depth, read length, and modification rate on clustering accuracy. Beta coefficients from the regression analysis were standardized to the same units to enable comparisons across features (larger beta coefficients correspond to a larger effect).

### Single molecule PORE-cupine (sm-PORE-cupine)

#### Preprocessing of raw signals

Nanopore signal data was preprocessed by performing z-score normalization on the raw signal before segmenting the traces into events using a change-point detection algorithm^26^ (from the Ruptures library). To reduce memory usage and compute time, we used discrete windows containing 5000 traces. In addition, we used a penalty of 2 to identify segmentation events more sensitively. The polyA adapter was identified based on its expected current and standard deviation signal from the kmer model and removed before further processing.

### Base-calling and mapping of nanopore reads

Reads were base-called with Guppy version 4.2.2+effbaf8. Both passed- and failed-quality reads from nanopore direct RNA sequencing were mapped to reference sequences using minimap2^13^ with the following parameters: -ax map-ont -F2064. For the 14 riboswitch and riboSNitch datasets, reads were mapped to the corresponding reference sequences. For SARS-CoV-2 data, reads were mapped to EPI_ISL_407987, as used by Kim et al^27^. For *C. albicans* transcriptome mapping, reference sequences obtained from the Candida Genome Database (CGD) were used^28^. For reads that were not successfully mapped by minimap2, their original fast5 files were subset out for mapping with DTW-based alignment.

### DTW-based signal alignment

For unmapped reads, subsequence Dynamic Time Warping (subseqDTW)^29^ was used to generate alignments using reference signals built from RNA kmer models (k=5). For riboswitch and riboSNitch data, subsequence Dynamic Time Warping (subseqDTW) was used for both mapping and signal alignment. Euclidean distances (similarity score) for a read signal when aligned against a reference signal (as computed by subseqDTW) were used to identify the best reference match for a read. The difference in scores between the best match and the next best match reference was computed as *Diff*_*Score* = 100 (*next*_*best*_*score* − *best*_*score*)/*next*_*best*_*score*, and a cutoff on this score was used to identify unique matches (cutoff determined based on evaluations that take minimap2 mapped reads as a standard of truth).

For accommodating mapping to the *C. albicans* transcriptome, a GPU-accelerated cuDTW stage was used while considering only the last 256 events of each unmapped read. An outlier removal stage was used to improve the robustness of mapping (normalized signal current > 3 standard deviations from the mean). The best reference identified by the cuDTW stage was then aligned using subseqDTW^29^ to align event signal levels with expected signal levels.

### Modification calling and BMM clustering

After aligning of read signals to reference signals, events corresponding to the same reference 5-mer were collapsed. We then used PORE-cupine^8^ to identify NAI-N3-induced RNA modifications along individual RNA strands. BMM clustering was then applied to separate RNA structure populations.

### Benchmarking analysis with riboswitch and riboSNitch data

To determine the accuracy of structure ensembles identified by sm-PORE-cupine, known structured riboswitches and riboSNitches were used for benchmarking. We performed structure probing of the riboswitches in the presence and absence of their ligands experimentally, as well as structure probing for wild-type (WT) and mutant (MT) versions of the riboSNitches. After nanopore sequencing and processing with the sm-PORE-cupine pipeline, we mixed no-ligand/ligand or WT/MT reads in equal proportion and assessed BMM clustering’s ability to accurately separate reads coming from the two libraries.

To test the sensitivity of structure cluster separation using BMM, we mixed reads at different proportions from the ADD No-ligand and Ligand libraries. The BMM was then used to cluster the mixtures; the proportion of No-ligand library obtained from BMM was defined as the predicted proportion and compared to the actual proportion (**Supplementary Figure 5g**).

### Analysis of heterogeneity for SARS-CoV-2 genome and the *C. albicans* transcriptome

#### Calculating a homogeneity score for each window

BMM clustering was restricted to two clusters for each transcript/window, even though potentially there can be more than two clusters. Based on BMM reported clusters for each window, we then calculated a homogeneity score that takes into account the degree of similarity between clusters and the relative proportions of the clusters. The homogeneity score serves as a weighted similarity score that is higher for structurally similar clusters and also higher in situations where there is a dominant cluster (as opposed to an even 50-50 split in proportions):

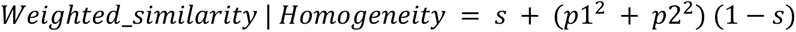

where *s* is the Pearson correlation between the two clusters and *p*1 and *p*2 are their respective proportions. It can be interpreted as an estimate of the expected similarity of two randomly sampled RNA structures for a transcript or window from its structure population, assuming that the two clusters and associated structures are correct.

#### Calculating heterogeneity on the SARS-CoV-2 genome

To identify heterogeneous windows along the SARS-CoV-2 genome, we used a 600nt window and a 10nt step size to scan through the entire length of the genome. We subsampled each window to 2000 reads before performing BMM clustering. To identify heterogeneous structures along specific sub-genomic RNAs, we first mapped the reads to full-length sub-genomic RNAs by ensuring that each read starts within the first 15 bases of the leader sequence and that the length of the read covers more than 85% of the full length of the sub-genomic RNA. We then performed BMM clustering on 600-nucleotide windows along each sub-genomic RNA. The homogeneity score was calculated for each window.

#### Calculating heterogeneity on the C. albicans transcriptome

For analysis of the *C. albicans* transcriptome, 100-nucleotide windows were used to scan through all transcripts. We subsampled each window to 2000 reads before performing BMM clustering for the libraries generated at 30°C and 37°C, respectively. The homogeneity score was calculated for each window.

### Analysis of *C. albicans* decay data

Raw reads were trimmed for adapter sequences and filtered for low-quality reads using the software Cutadapt^30^. The trimmed reads were mapped to the *C. albicans* transcriptome using Bowtie2 with default parameters^31^. The raw counts of each transcript were quantified using featureCounts^32^. FPKM values for each gene were calculated across different timepoints post structure probing (0mins, 20mins, 40mins, 60mins, 120mins). The FPKM values across 5 timepoints and at 30°C and 37°C respectively, were analysed using StepMiner^20^ to determine significantly stable and decaying transcripts. StepMiner performs an adaptive regression that identifies step-wise transitions in a time-series data, which gives insights into the timing of the gene expression-switching event.

### Modelling of RNA secondary structures

RNA secondary structures based on BMM predicted clusters were modelled using the ‘RNAstructure’^33^ tools ‘partition-smp’ and ‘stochastic’, using the aggregate reactivity of the cluster as partition functions to constrain the secondary structure model. The centroid of the output from ‘stochastic’ was calculated and visualized using VARNAv3-93.

### Statistical and visualization analysis

The processing, analysis, and visualization of the data were performed using scripts with python-3.9.6, and the associated modules snakemake −7.15.2, pyfaidx-0.6.1, scipy −1.7.1, tslearn-0.5.2, h5py-3.8.0, matplotlib-3.5.1, numpy-1.21.5, pandas-1.5.0, ruptures-1.1.6, seaborn-0.11.2, intervaltree-3.1.0, and scikit-learn-0.24.2.

## Data availability

We are currently submitting the sequencing data to the Gene Expression Omnibus (GEO) database, and we will update the link here when the submission is done.

## Code availability

The source code for all analyses in this manuscript is freely available from GitHub under a GPLv3 license (https://github.com/YueLab-GIS-ASTAR/sigpore). Downstream analysis scripts can be found in the folder ‘analysis/notebooks’.

## Acknowledgements

We thank members of the Wan lab, Yue Wang lab and Niranjan lab for their helpful discussions. Y.W. is supported by funding from A*STAR Investigatorship, National Research Foundation of Singapore, the European Molecular Biology Organization Young Investigatorship, and the Canadian Institute for Advanced Research Azrieli Global Scholar fellowship.

## Author contributions

Y.W. and J.W. conceived the project. Y.W., J.W. and Y. Wang designed the experiments. J.W., W.T.T., and G.Z performed all the experiments. N.N. supervised the analysis. J.H, A.A. and performed all the computational analysis. Y.W, J.W. J.H. and N.N. organized and wrote the paper with all other authors.

## Supplementary Figure legends

**Supplementary Figure 1.**
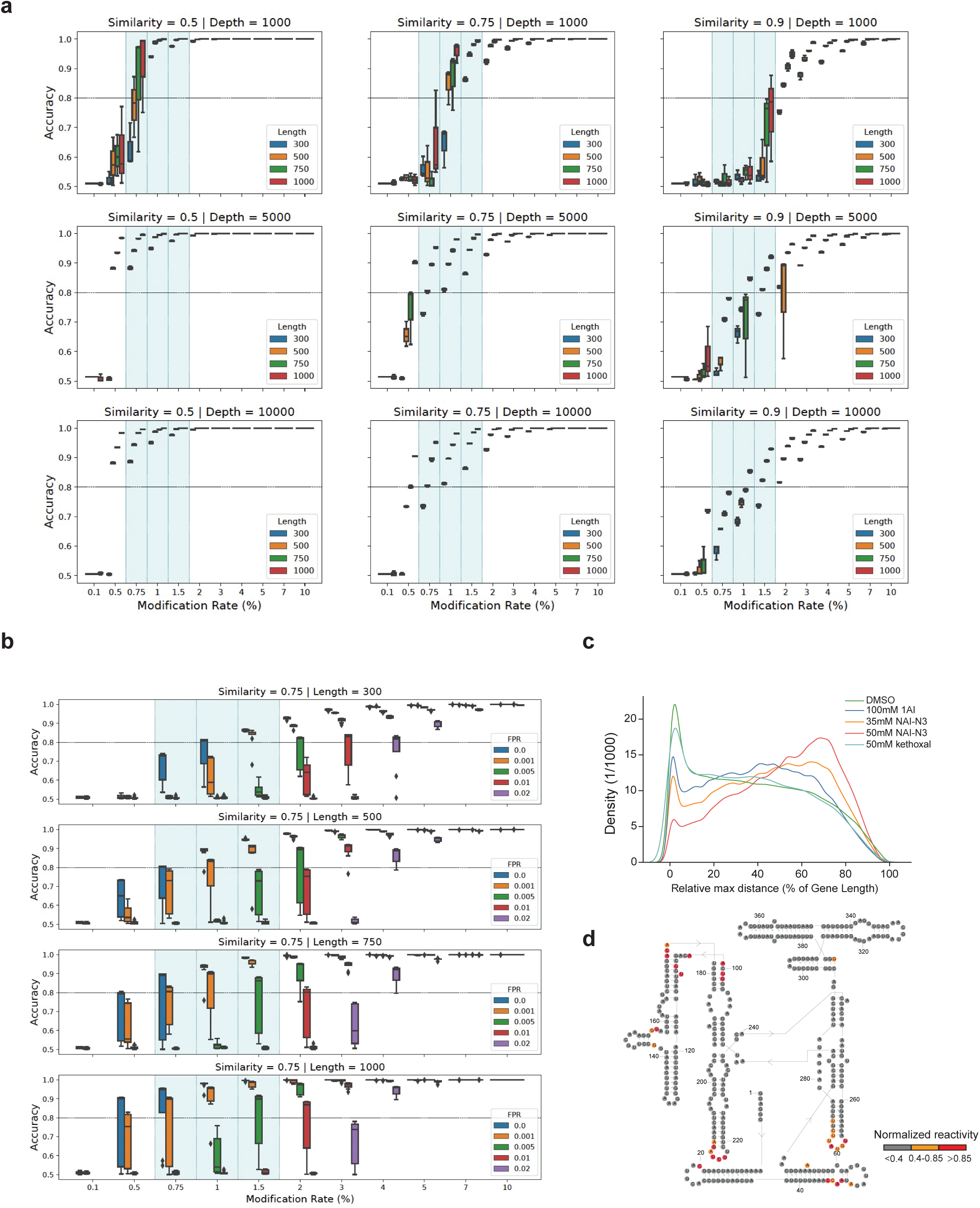
Identification of key parameters that are important for accurate clustering. **a,** Boxplots showing the distribution of clustering accuracies at different structural similarities, sequencing depths, transcript length, and modification rates for simulated transcripts using BMM clustering (Methods). **b**, Boxplots showing the distribution of clustering accuracies at different false positive rates (for identifying single-stranded bases), transcript lengths, and modification rates for simulated transcripts using BMM clustering (Methods). **c,** Line plot showing the kernel-density estimated distribution of all distances between the furthest 2 modified positions on a read, for the different probing conditions tested. **d**, NAI-N3 reactivity is determined using PORE-cupine and was mapped to the known secondary structure of the Tetrahymena ribozyme. Highly, moderately, and poorly reactive bases are coloured in red, orange, and grey, respectively.

**Supplementary Figure 2.**
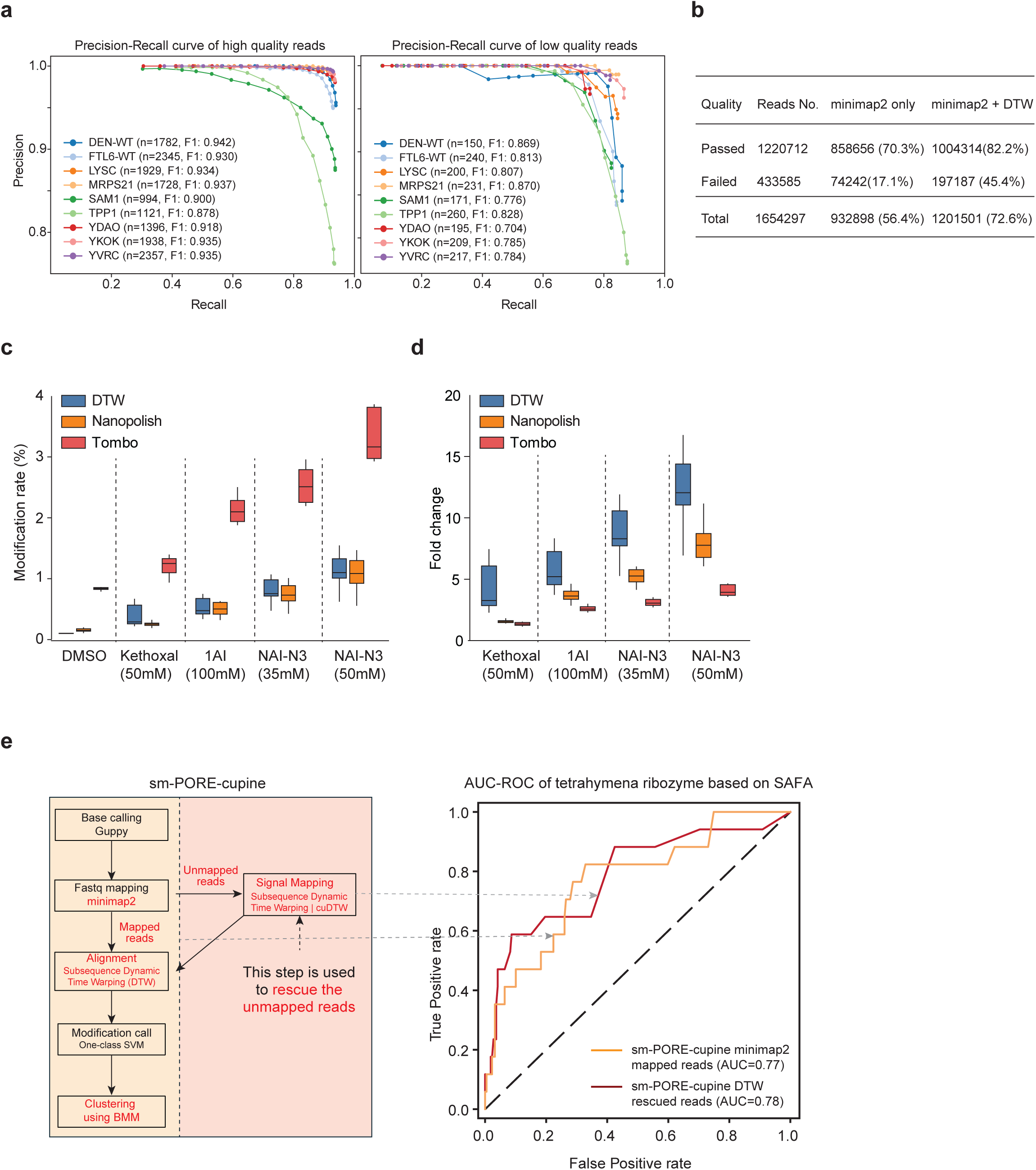
Structure information obtained using sm-PORE-cupine is sensitive and accurate. **a**, Precision-Recall curves for subsequence Dynamic Time Warping (subseqDTW) alignments for high-quality direct RNA sequencing reads (Quality>=7, ***Left***) and low-quality sequencing reads (Quality<7, ***Right***), using minimap2 alignments as a truth set. **b**, Table showing the percentage of reads that pass (Quality>=7) or fail (Quality<7) quality filters and can be mapped by minimap2 or minimap2 combined with DTW. **c**, Boxplot showing the modification rate detected for RNAs treated with different chemical compounds, using different re-squiggling methods. Both Nanopolish and Tombo use minimap2 for sequence mapping before re-squiggling. **d**, Boxplot showing the fold enrichment of modification rate over false positives obtained on RNAs treated with different chemical compounds, using different re-squiggling methods. **e, *Left***, schematic showing the computational workflow of sm-PORE-cupine. AUC-ROC curve of the sm-PORE-cupine minimap2 mapped reads (orange) and sm-PORE-cupine DTW rescued reads (red) based on the known secondary structure of the Tetrahymena ribozyme. ***Right***, line plots showing the AUC-ROC for the minimap2-mapped reads (orange) and DTW-rescued reads (red).

**Supplementary Figure 3.**
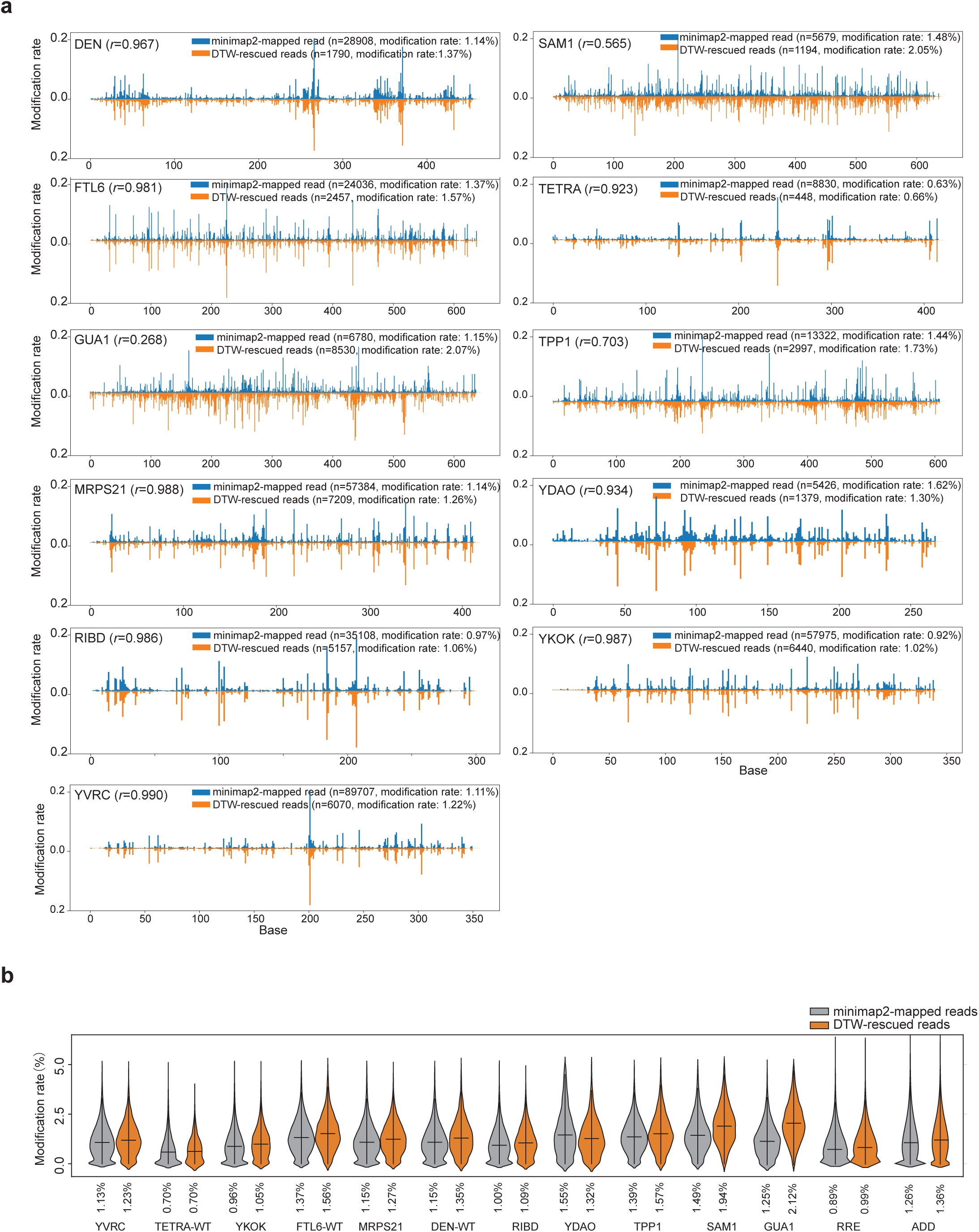
DTW-rescued reads have a similar reactivity pattern and a higher modification rate than minimap2-mapped reads. **a,** Line plots showing the modification rates along the different RNAs when the reads are mapped using minimap2 (blue) or DTW (orange). Pearson correlation is used to calculate the similarity in reactivity between minimap2-mapped and DTW-rescued reads. **b**, Violin plots showing the distribution of modification rates in reads that are mapped by minimap2 versus rescued by DTW.

**Supplementary Figure 4.**
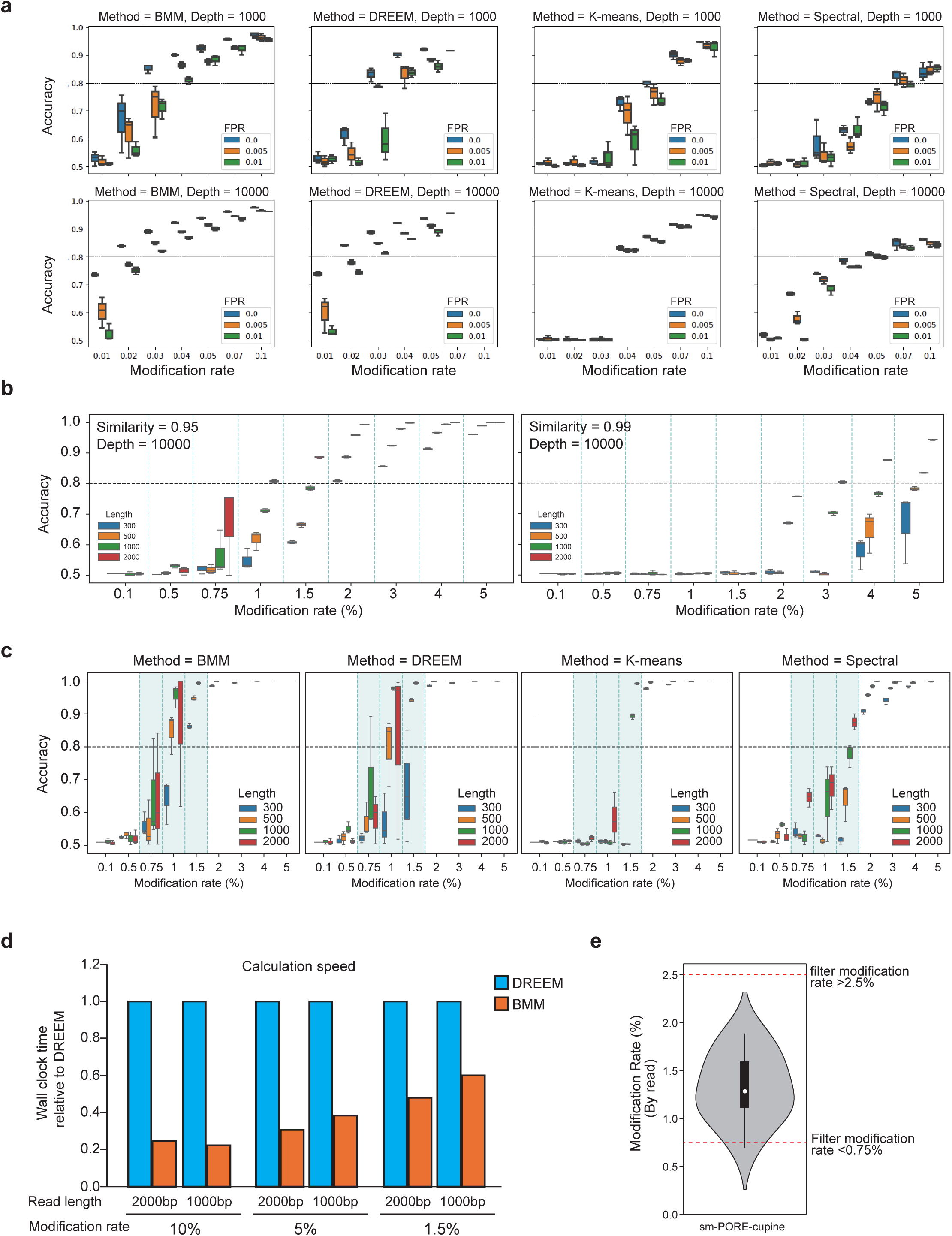
Identification of the clustering strategy for accurate clustering of RNA structure populations using simulation. **a,** Boxplot showing the distribution of clustering accuracies using different clustering strategies, at different sequencing depths, modification rates, and false positive rates (FPR). Mixture models inferred with EM algorithms are more robust to low modification rates. Results of all clustering methods are highly sensitive to errors, especially at lower levels of modifications (∼1%). DREEM failed to converge at a 10% modification rate. **b**, Boxplots showing the distribution of clustering accuracies for simulated transcripts, with different lengths, with 95% (left) and 99% (right) similarities in reactivity. **c**, Boxplots showing the distribution of clustering accuracies using different clustering strategies on simulated transcripts with different lengths and modification rates. **d**, Barplots showing the calculation speed using DREEM and BMM for different read lengths and modification rates. **e**, Violin plot showing the distribution of the modification rates of all sequenced reads. The red dashed lines indicate the thresholds for lowly modified reads (25% of total reads) and extremely highly modified outlier reads.

**Supplementary Figure 5.**
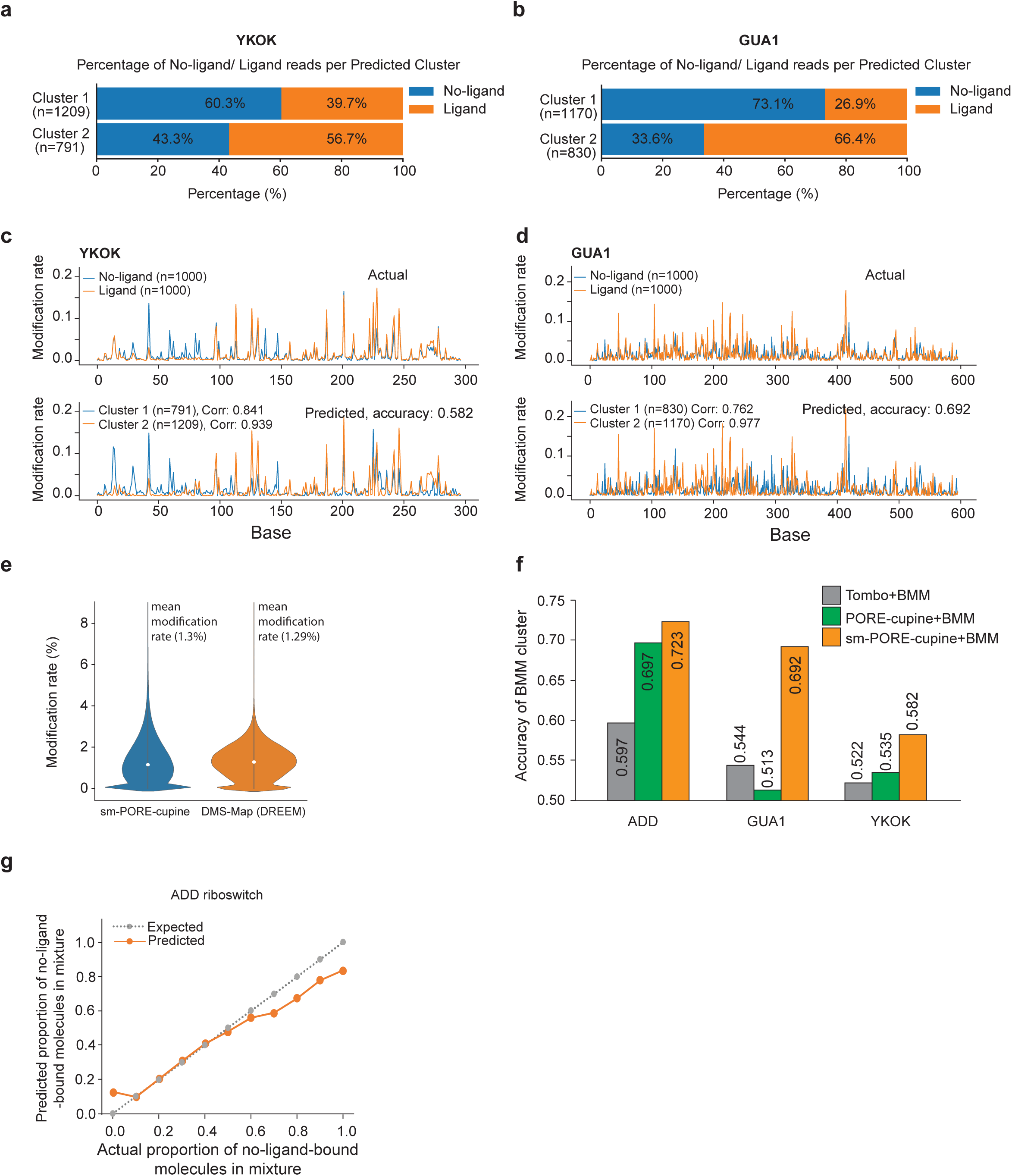
sm-PORE-cupine can separate Ligand-bound and non-ligand-bound riboswitches. **a, b,** Barcharts showing the assignment of reads from Ligand-bound and No-ligand-bound YKOK (**a**) and GUA1(**b**) riboswitch into two structure populations by sm-PORE-cupine. **c, d**, ***Top***, Line plots showing the aggregate reactivity of the YKOK(**c**) and GUA1(**d**) riboswitch when it is structure probed in the absence (blue) or presence (orange) of their ligands. ***Bottom***, Line plots showing the aggregate reactivity of the two clusters determined using BMM clustering for the YKOK(**c**) and GUA1(**d**) riboswitch. **e**, Violin plots show the distribution of modification rates of the reads present in sm-PORE-cupine and in the published DMS-MaP-DREEM pipeline. **f**, Barplots show the accuracy of BMM clusters for 3 riboswitches using different pipelines as shown. **g**, Line plot showing the predicted versus actual proportion of No-ligand-bound molecules using BMM when different proportions of Ligand and No-ligand-bound ADD riboswitch molecules are mixed together.

**Supplementary Figure 6.**
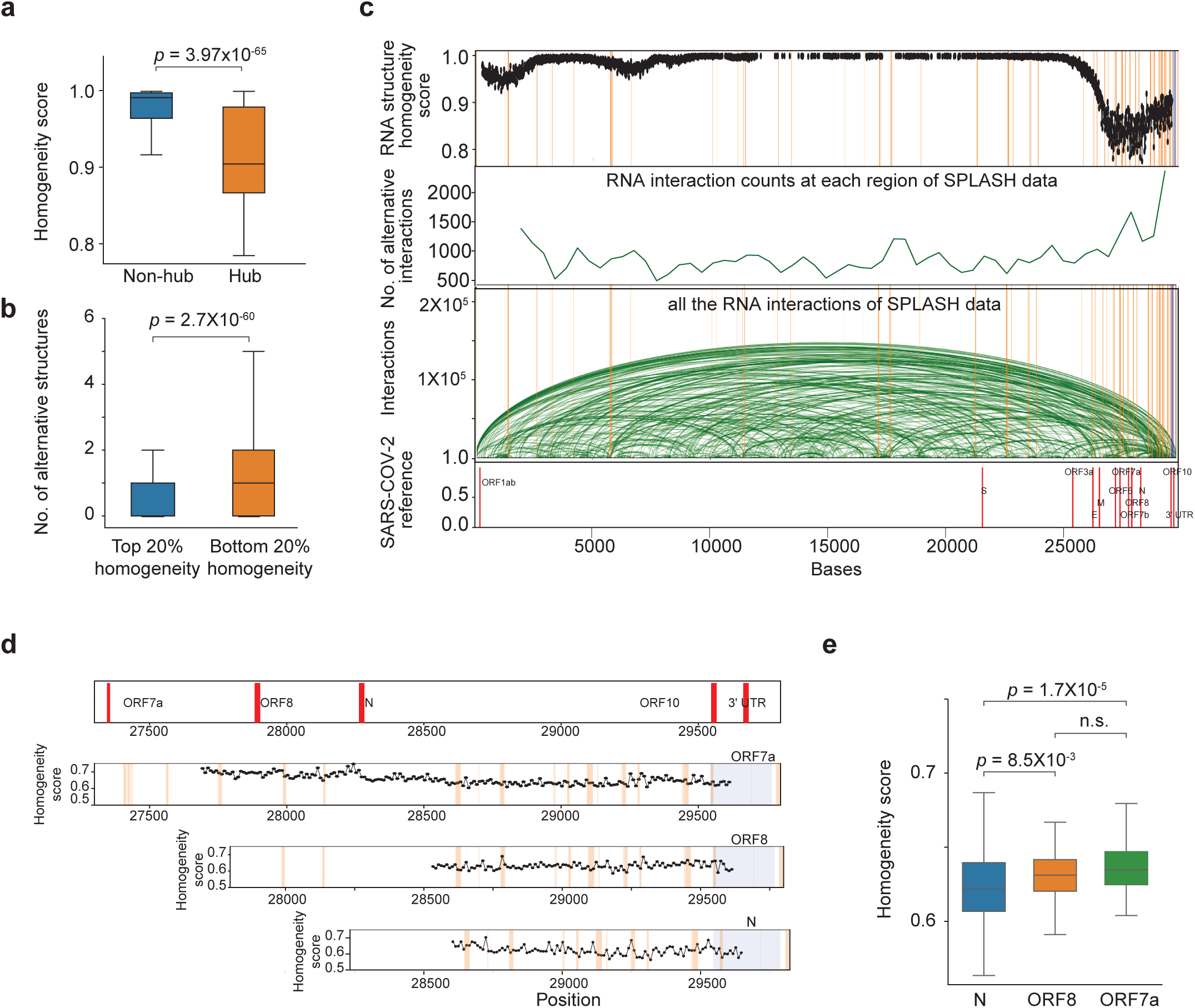
sm-PORE-cupine can identify RNA structure ensembles from SARS-CoV-2 subgenomic RNAs. **a,** Box plots showing the homogeneity scores of non-hub and hub regions along SARS-CoV-2. Regions with the highest 5% of alternative interactions by SPLASH were defined as hubs. *p*-values are calculated using Wilcoxson Rank Sum test. **b**, Boxplots showing the distribution of alternative SPLASH interactions for the top and bottom 20% homogeneous regions along the SARS-CoV-2 genome. *P*-values are calculated using Wilcoxson Rank Sum test. **c, Top**, Line plot showing the degree of RNA structure homogeneity along the SARS-CoV-2 genome. **Middle,** Lineplot showing of frequency of alternative RNA-RNA interaction data from our previous Splash data^17^. High frequencies indicate that the position interacts with many other positions, suggesting highly complex alternative structures. **Bottom,** Arc plots showing pair-wise RNA-RNA interactions based on our published SPLASH data^17^, along the SARS-CoV-2 genome. The orange lines indicate the top 5% locations with the highest number of alternative RNA-RNA interactions found in SPLASH. **d**, Line plots showing the homogeneity score at each window in each subgenomic RNAs ORF7a, ORF8, and N. The orange lines indicate the top 5% locations with the highest number of alternative RNA-RNA interactions found in SPLASH. **e**, Boxplots showing the homogeneity scores on sgRNA ORF7a, 8, and N. *P*-values are calculated using the Wilcoxon Rank Sum test.

**Supplementary Figure 7.**
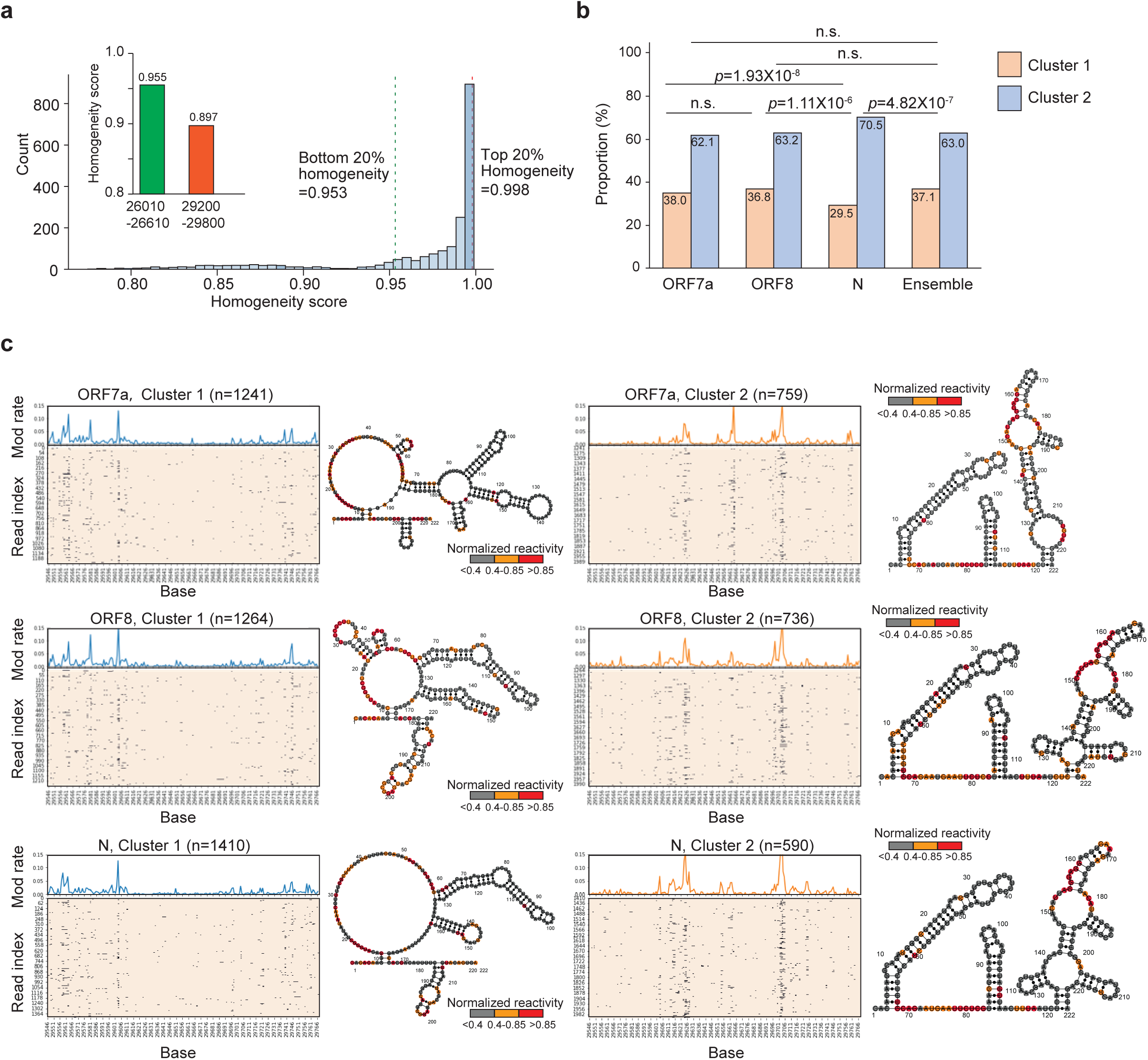
A 3’UTR RNA region in the SARS-CoV-2 genome forms different proportions of structure ensembles in subgenomic RNAs. **a**, Histogram showing the distribution of homogeneity scores along the SARS-CoV-2 genome. The dotted lines indicate the top (red) and bottom (blue) 20% of the homogeneity scores. The inset shows the homogeneity score of a 3’UTR region and another independent window upstream. **b**, Barplots showing the proportions of the two clusters in the different subgenomic RNAs, including ORF7a, ORF8, N, and when the RNAs are aggregated together and not separated into individual sgRNA (ensemble). *P*-values are calculated using the chi-square test of association. **c**, ***Left***, For each sgRNA, we show a heatmap that indicates the distribution of NAI-N3 modification along each read in the population. We show the aggregated RNA structure signal as a line plot at the top. The aggregate structure signals of the two populations are in blue (Cluster 1) and in orange (Cluster 2), respectively. ***Right***, we performed RNA structure modelling using the aggregate reactivity in each cluster as constraints. Highly, moderately, and poorly reactive bases are coloured in red, orange, and grey, respectively.

**Supplementary Figure 8.**
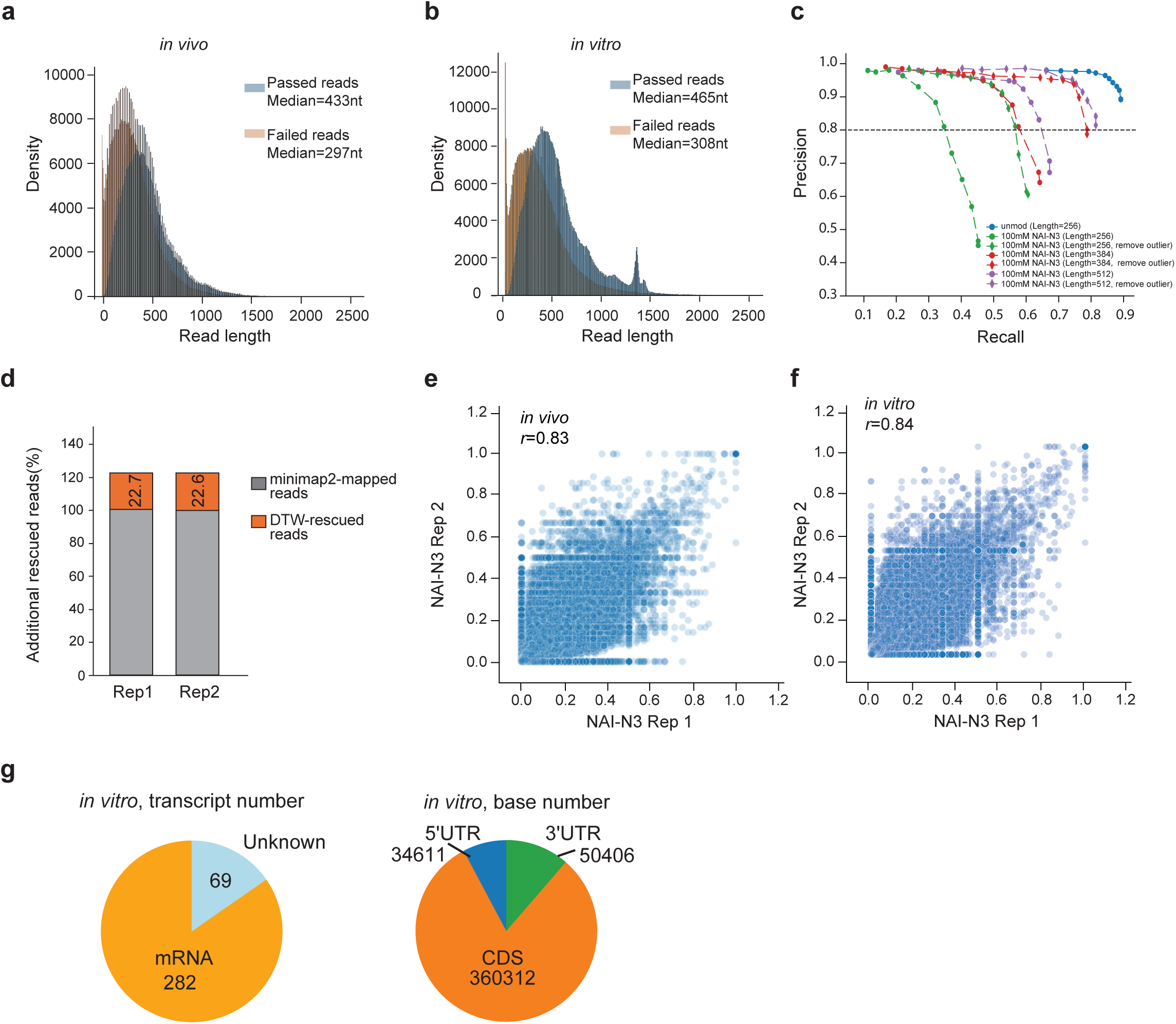
Features of the RNA structure landscape in *C. albicans*. **a, b**, Density plot showing the distribution of read length from direct RNA sequencing for passed-quality and failed-quality reads for both *in vivo* (**a**) and *in vitro* (**b**) samples. **c,** Precision-Recall curves showing the precision and recall of signal using subsequence Dynamic Time Warping (cuDTW) on the *C. albicans* transcriptome. We have tested different thresholds, including length (the last 256nt, 384nt, and 512nt of each read), before or after filtering the outliers based on normalized Z-score values, and the normalized Z-score values beyond three standard deviations of its mean are defined as outliers. This data shows that the Precision-Recall was improved after filtering out outliers. The subseqDTW-based analysis for the last 256nt is used to accelerate transcriptome-wide mapping with a relatively decent precision. **d**, Bar charts showing the percentage of reads that can be mapped for each RNA in the *C. albicans* transcriptome, using either minimap2 alone or minimap2 with DTW. The percentage of additional reads rescued when applying subseqDTW on reads that failed to be aligned by minimap2 is in orange. **e, f**, Scatter plots showing the correlation of RNA structure reactivity between biological replicates *in vivo* (**e**) and *in vitro* (**f**). Each dot is a base. **g**, Pie chart showing the number of detected transcripts (***Left***) and bases (***Right***) in *C. albicans* RNA *in vitro* using sm-PORE-cupine.

**Supplementary Figure 9.**
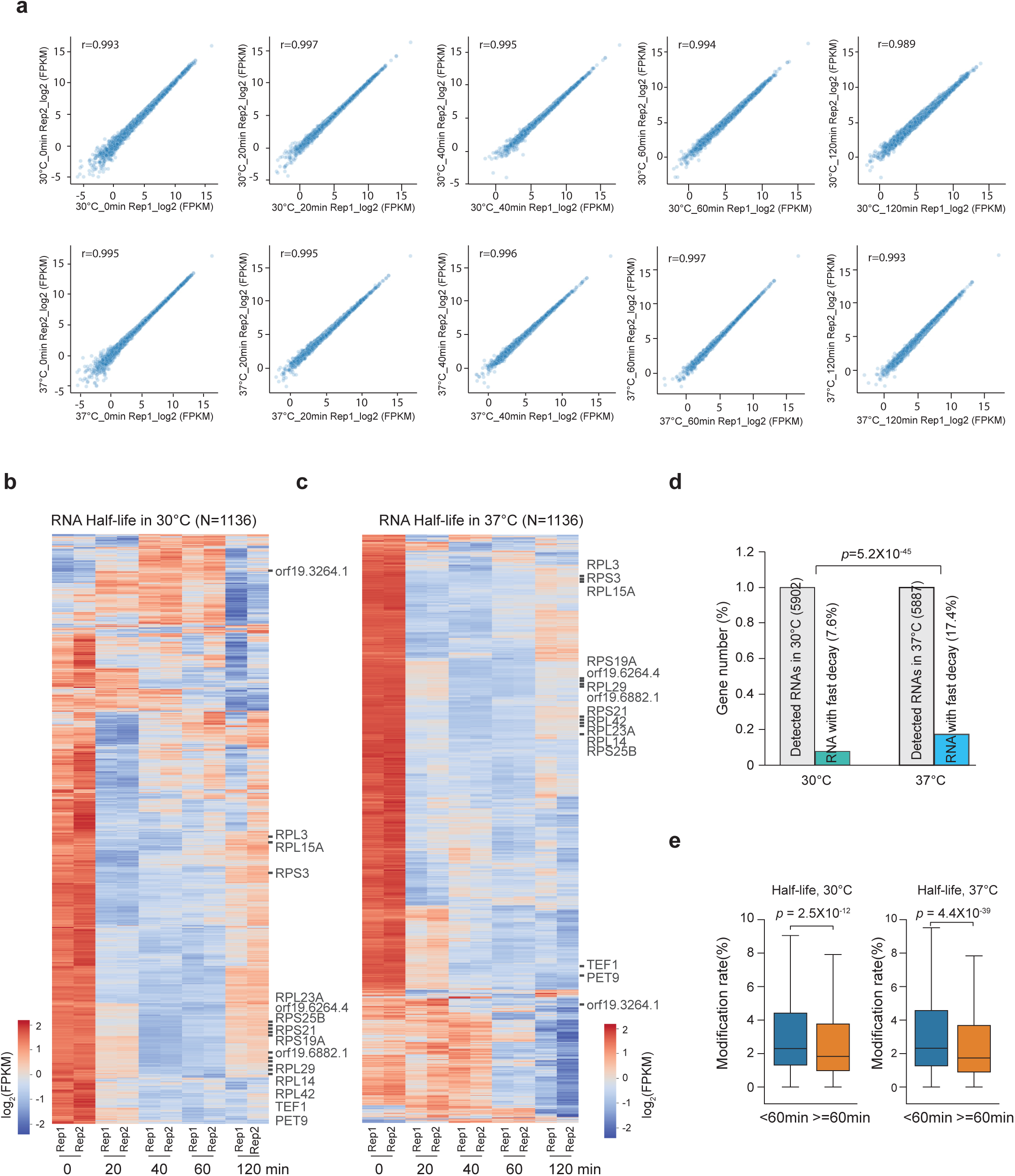
RNA decay measurements in *C. albicans* are reproducible. **a,** Scatterplots showing the correlation between two biological replicates of *C. albicans* RNA sequencing data at 0, 20, 40, 60, and 120min after treating the cells with Thiolutin at 30°C and 37°C. **b, c,** Heatmap, generated using Stepminer (**Methods**), showing the expression level (log_2_ FPKM) of C. albicans transcriptome at 0, 20, 40, 60 and 120min after treating the cells with Thiolutin at 30°C (**b**) and 37°C (**c**). **d**, Barplots showing the number of transcripts that are degraded rapidly at 30°C and 37°C. *P*-values are calculated using the chi-square test of association. **e**, Boxplot showing the distribution of RNA modification rates in transcripts that are degraded rapidly versus slowly. *P*-values are calculated using the Wilcoxon Rank Sum test.

**Supplementary Figure 10.**
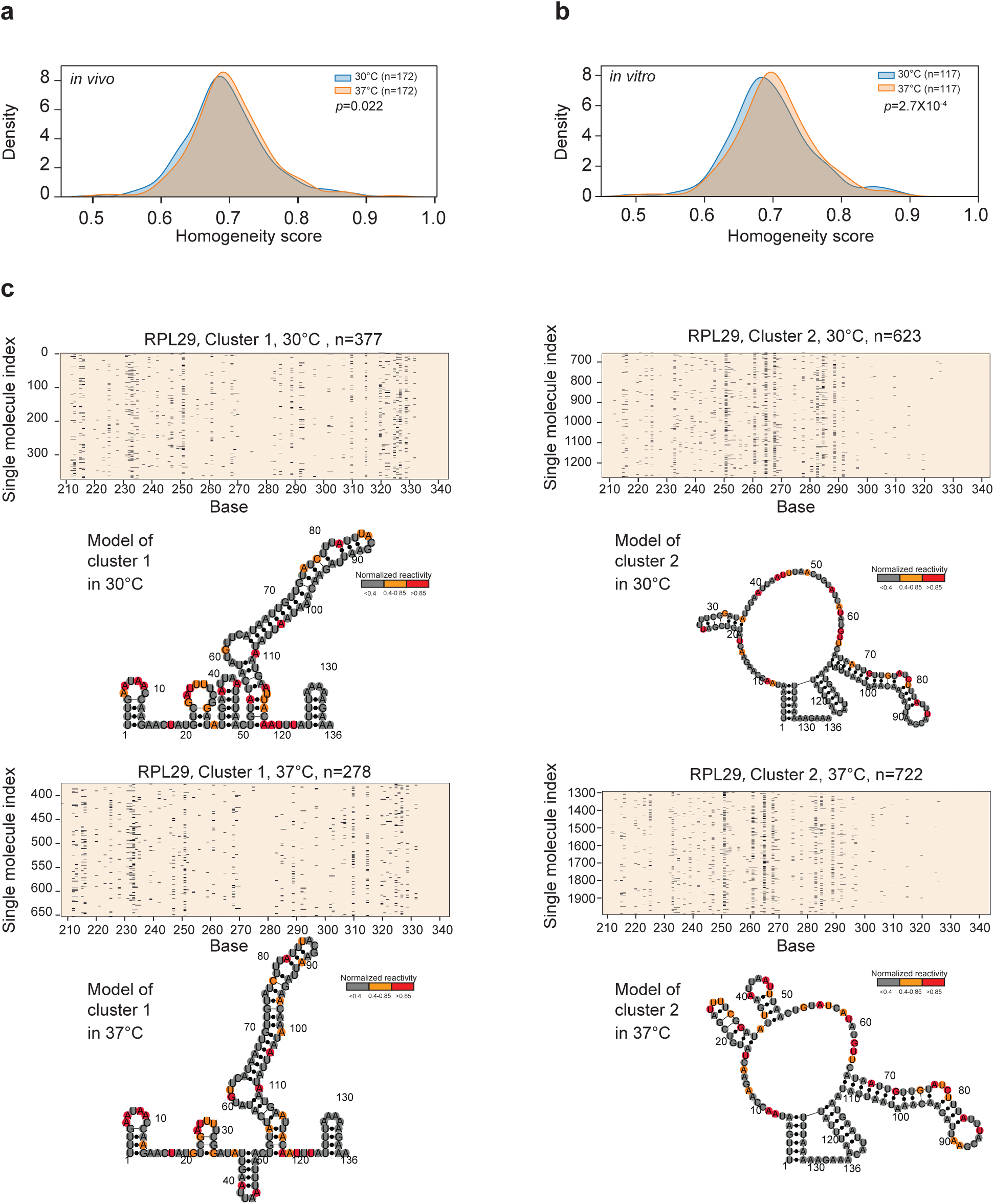
Structure modelling of RNA ensembles for test genes. **a, b**, Density plots show the distribution of homogeneity scores in the *C. albicans* transcriptome, at 30°C and 37°C, *in vivo* (**a**) and *in vitro* (**b**). *P*-values are calculated using the two-sample Kolmogorov-Smirnov test. **c**, ***Top***, Heatmap showing the distribution of NAI-N3 modifications along each molecule in each cluster of RPL29 using BMM clustering. ***Bottom***, RNA structure modelling is performed using the aggregate reactivity of each cluster as constraints. Highly, moderatel,y and poorly reactive bases are coloured in red, orange, and grey, respectively.

**Supplementary Figure 11.**
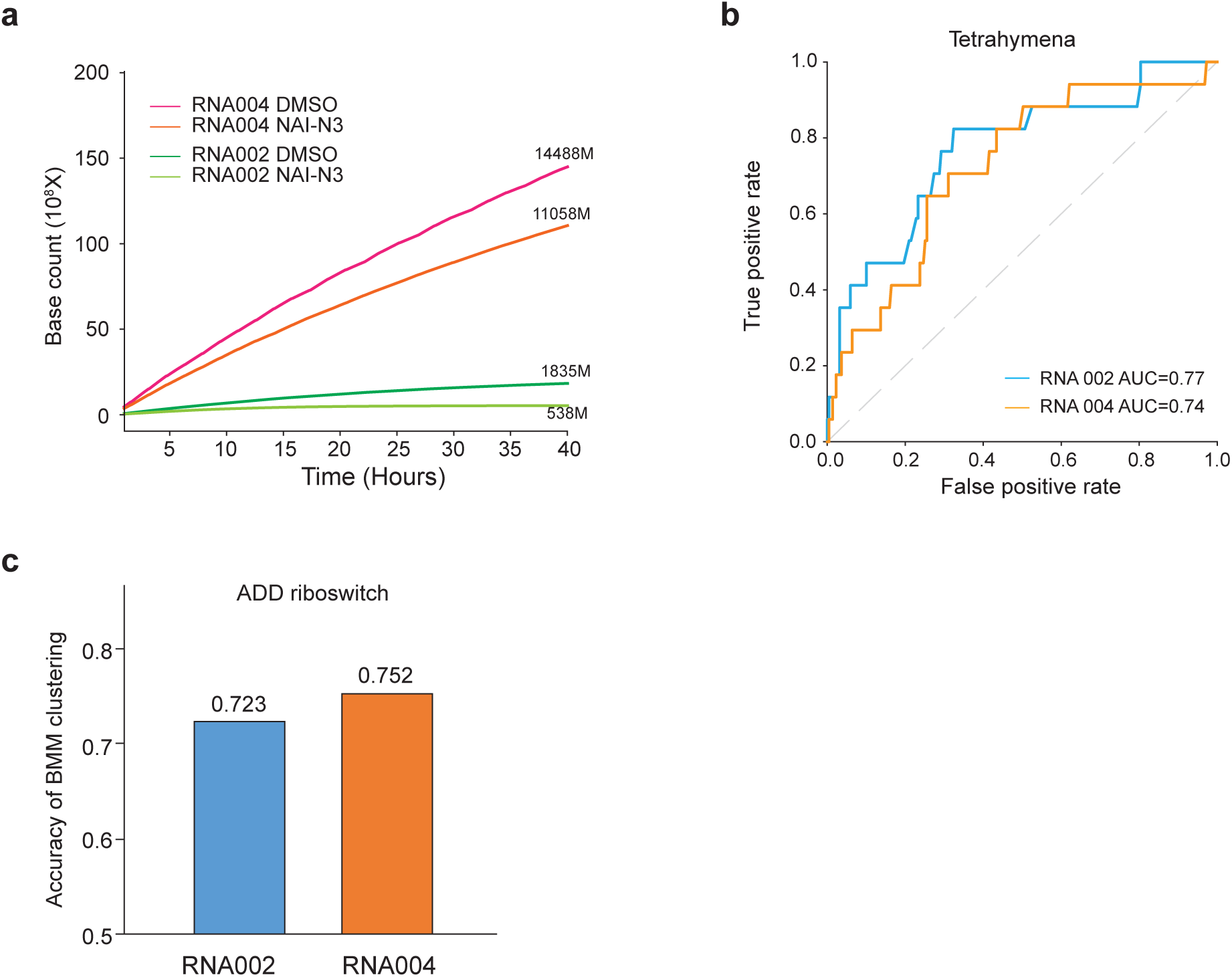
RNA 004 kit can be used to determine RNA structure ensembles. **a,** Line plots showing the sequencing depth (Base count) for DMSO and NAI-N3-treated RNAs at different times (hours) using the direct RNA sequencing RNA002 and RNA004 kit, respectively. **b**, AUC-ROC were calculated using the reads from the RNA002 and RNA004 kits, based on the experimental footprint (SAFA) of the Tetrahymena ribozyme. **c**, Barplots showing the cluster separation accuracy of ADD riboswitches using sm-PORE-cupine, using reads prepared from the RNA002 and RNA004 kit, respectively.

